# Notch1 induces endothelial plasticity to mediate hyaloid vessel involution

**DOI:** 10.1101/2025.10.01.679581

**Authors:** Khazaal Shaymaa, Sango Abdoul-Razak, Megne Ariane Vanessa Megne, Silva Sosa Alexa, Zouine Kawtar, Gaelle Mawambo, Chidiac Rony, Bora Kiran, Chen Jing, Oubaha Malika

## Abstract

Hyaloid vascular regression is a critical developmental process essential for vitreous transparency and normal vision, yet the molecular cues orchestrating its involution remain incompletely defined. Here, we identify Notch1 as a pivotal regulator of hyaloid vessel clearance, acting independently of apoptosis to coordinate endothelial detachment, transient plasticity, and migration.

Using an endothelial-specific Notch1 knockout mouse model, we demonstrate that loss of Notch1 results in persistent hyaloid vasculature characterized by excessive proliferation and stabilization of the vascular network. Mechanistically, Notch1 activation during the regression window induces endothelial-to-mesenchymal transition (EndoMT) marked by *Snail1* and *Slug* upregulation. This transcriptional signature is accompanied by detachment of endothelial cells from the vascular tubes. In contrast, Notch1-deficient hyaloid vessels retain endothelial cells stably adherent to the vessel wall. Further analysis reveals that Wnt receptors FZD4, LRP5 and LRP6 previously implicated in hyaloid involution are transcriptionally downregulated in Notch1-deficient hyaloids, suggesting that the collaboration between these processes may occur through crosstalk between the Notch and Wnt pathways.

Collectively, our findings uncover a Notch1-driven multicellular regression program that governs developmental vessel regression, redefining the molecular principles of vascular pruning. These results have broad implications for understanding vascular remodeling in both physiological and pathological contexts and may guide therapeutic strategies to modulate vascular regression in ocular disorders.

**One-Sentence Summary:** Notch1 drives hyaloid regression through a multicellular program that defines an apoptosis-non-exclusive paradigm of vessel pruning.

## Introduction

Ocular vascular development is crucial for eye growth and function [1]. Three vascular networks sustain the eye: the retinal, choroidal and transient hyaloid vessels. The hyaloid system provides oxygen and nutrients to the developing lens and inner retina but undergoes a tightly timed regression as the retinal vasculature expands. In mice, hyaloid vessels regress shortly after birth and are largely eliminated by postnatal day 16-21. In humans, regression begins in late gestation (weeks 28-30) coinciding with the maturation of the retinal circulation [1–4].

Hyaloid vessels are composed of three principal cellular constituents including endothelial cells, pericytes, and hyalocytes, a specialized macrophage-like cell population residing in the vitreous. Endothelial cells form the inner lining of hyaloid vessels and are pivotal in the regression process during ocular development. They do not only contribute structurally but also actively regulate the vascular remodeling through tightly controlled processes of proliferation, apoptosis, and interaction with pericytes and macrophages [5–7]. During hyaloid vessel regression, endothelial cells undergo apoptosis, a process influenced by signals such as vascular endothelial growth factor (VEGF) downregulation and macrophage-mediated phagocytosis, leading to the dismantling of the vessel network. These coordinated actions ensure proper ocular maturation and the clearance of transient hyaloid vessels [2, 5, 8–10]. Pericytes, closely associated with endothelial cells, maintain vascular integrity through basement membrane deposition and PDGF-β/PDGFRβ and TGF-β signaling, whereas their loss during hyaloid regression destabilizes endothelial cells and drives vessel involution [11–13]. Hyalocytes, orchestrate immune surveillance and vascular pruning, acting as key effectors of hyaloid regression [5, 8]. Together, these cell populations integrate cues from different signaling pathways to guide the dynamic formation and regression of the hyaloid vasculature during ocular development. Failure of hyaloid vascular regression is associated with a spectrum of congenital disorders characterized by aberrant ocular vascularization and disrupted retinal development, often culminating in vision loss or blindness. These include Persistent Fetal Vasculature (PFV), Norrie disease, retinopathy of prematurity (ROP), familial exudative vitreoretinopathy, and Coats disease [14–16]. The etiology of these disorders frequently involves mutations in genes regulating developmental angiogenesis, resulting in the pathological persistence of the hyaloid vasculature. PFV, a rare congenital anomaly, arises from involution failure of hyaloid vessels and is often associated with mutations affecting Wnt signaling components, including Wnt ligands and Frizzled receptors [5, 17, 18]. Similarly, mutations in the *NDP* gene underlie Norrie disease, an X-linked disorder causing severe retinal dysgenesis in males [19]. In contrast, ROP stems from delayed and abnormal retinal vascularization in preterm infants and is frequently accompanied by persistent hyaloid remnants. Major risk factors of ROP include extreme prematurity and fluctuations in oxygen tension during neonatal care, which destabilize VEGF-mediated vascular growth [20–22]. While no definitive cures exist, clinical management includes vitrectomy to address structural complications of PFV, and laser photocoagulation or anti-VEGF therapy to limit neovascularization in ROP [22–24]. Emerging gene therapy strategies targeting *NDP* and other angiogenic regulators are currently under investigation as potential avenues for restoring retinal structure and function [25]. Further investigation into the molecular pathways underlying vascular regression and retinal development is essential, as a deeper understanding of these processes may lead to the identification of novel therapeutic strategies. Elucidating the molecular mechanisms that govern hyaloid vascular regression, and retinal vascular development is fundamental to understanding normal ocular morphogenesis and to identifying therapeutic entry points for congenital and neovascular retinal diseases. Beyond the traditionally emphasized role of apoptosis [2, 6], research study highlights the importance of coordinated vascular remodeling, signaling crosstalk, and endothelial cell cycle regulation in mediating hyaloid vessel involution. The Wnt ligand Wnt7b, secreted by macrophages closely associated with hyaloid vessels, has been shown to initiate canonical Wnt/β-catenin signaling in neighboring endothelial cells through interaction with Frizzled-4 (FZD4) and LRP5 receptors [5]. This signaling cascade contributes not only to the destabilization of the vessel wall but also to the modulation of endothelial cell behavior, including cell cycle arrest and loss of proliferative competence which are key hallmarks of vascular regression. *In vivo* genetic studies further support this model. Conditional deletion of the pro-apoptotic regulator Bcl-2 interacting mediator of cell death (Bim) in endothelial and mural cells attenuates hyaloid regression, but intriguingly, this phenotype is also associated with aberrant vessel remodeling and persistence of immature vascular structures [6]. Vascular regression involves an interplay of endothelial cell proliferation, matrix remodeling, and programmed cell death [26]. While Wnt signaling has been implicated, Notch pathways well established in retinal angiogenesis and vascular stabilization [27–32], remain unexplored in hyaloid regression, though their roles in related remodeling processes point to potential regulatory functions. The Notch signaling pathway is a fundamental regulator of vascular development, playing a pivotal role in maintaining endothelial quiescence and limiting excessive vessel proliferation to preserve vascular homeostasis [33–35]. Given the centrality of both Wnt and Notch pathways in modulating endothelial cell fate decisions and vascular patterning, their coordinated activity likely orchestrates the delicate balance between angiogenesis, vessel maturation, and regression. Disruption of this balance may underlie aberrant ocular vascular phenotypes, such as PFV and related persistence disorders. Notch signaling, through its conserved mechanisms of cell-cell communication, dictates fate specification and differentiation across tissues, with pivotal roles in endothelial patterning and vascular morphogenesis. Among its ligands, members of the Delta-like ligand (DLL1, DLL3, DLL4) and the Jagged (JAG1, JAG2) families, DLL4 is uniquely and predominantly expressed by endothelial cells and serves as a critical determinant of arterial specification and branching morphogenesis [36, 37].. In the retina, Lobov et al. demonstrated that the activation of DLL4/Notch1 signaling facilitates vessel pruning and regression by modulating endothelial transcriptional programs toward a vasoconstrictive, anti-angiogenic state. The inhibition of DLL4 was found to reduce the extent of physiological regression in the developing retinal vasculature, suggesting a broader role for Notch1 in post-angiogenic remodeling and homeostatic vessel elimination [38]. While Notch1 activity in retinal vascular development is increasingly appreciated, its specific contribution to hyaloid vessel regression remains to be directly defined. Here, we investigate the cellular dynamics underlining hyaloid vascular regression, with a focus on the coordinated contribution of hyalocytes, pericytes, and endothelial cells. We identify a previously unrecognized role for Notch1 in hyaloid vessel pruning, where it controls endothelial proliferation and plasticity. Using inducible ubiquitous (*Notch1^fl/fl^; Cdh5-CreERT2*) and endothelial-specific (*Notch1^fl/fl^; CreESR1*) deletion models, we delineate the impact of Notch1 loss on hyaloid regression. Our findings show that hyaloid involution relies not only on apoptotic clearance but also on integrated signaling cues that shape endothelial fate. This work establishes a refined model of postnatal ocular vascular regression and highlights new strategies to tackle pathological vessel persistence in eye disease.

## Results

### Cells relocation, not apoptosis, drives hyaloid pruning

While apoptosis has been implicated in pruning redundant vasculature, its temporal and functional relevance to hyaloid vessel involution remains incompletely defined. In mice, the postnatal regression of hyaloid vessels occurs concomitantly with the growth of retinal vasculature (Fig. 1A). To explore the cellular basis of hyaloid regression, we delineated the developmental trajectory and early regression of the hyaloid vasculature in C57BL/6J mice, spanning late embryogenesis (E18.5) to early postnatal stages (P8). Isolectin B4 (IB4) labeling of flatmounted hyaloids revealed a progressive decrease in hyaloid vascular density over time (Fig. 1B). Quantification revealed a progressive decline, with ∼20% reduction by P0, ∼40% by P4, and ∼70% by P8 compared to E18.5. (Fig. 1C). In parallel, we observed a marked narrowing of vessels diameter (Fig. 1D) and a significant decrease in branching complexity relative to E18,5, as reflected by reduced vascular junctions (Fig. 1E). Previous reports have identified hyaloid vessels apoptosis at mid-gestation stages in the embryo; preceding the onset of postnatal vessel regression [2, 6]. However, a systematic longitudinal analysis has been lacking. To address this, we performed immunolabeling for cleaved caspase-3 (C-Casp3), a marker of apoptotic cells, across the regression timeline. Surprisingly, the frequency of apoptotic cells per 100µm of hyaloid vessels length remained constant from P0 through P8 (Fig. 1F, G), indicating that regression proceeds independently of increased apoptosis. Thus, we asked whether changes in endothelial cell proliferation might contribute to hyaloid involution. Immunostaining for the proliferative marker Ki-67 revealed a significant decline in proliferating cells within the hyaloid vasculature at P4 and P8 compared to P0 (Fig. 1H, I). Intriguingly, we observed a concomitant increase in proliferating cells located outside the hyaloid vascular structure. At P0, Ki-67^positive^ cells were predominantly intravascular, whereas by P8, the majority were extravascular (Fig. 1H, arrowheads). Quantitative analysis confirmed a 3- to 5-fold increase in detached proliferating vascular (endothelial or pericytes) cells per 100 µm in extravascular regions by P4 and P8, respectively (Fig. 1J). Together, these findings show that hyaloid vascular regression in mice occurs in the absence of increased apoptosis during postnatal stages. Instead, the regression is accompanied by a decline in vascular cell proliferation and a spatial shift in proliferative activity to the extravascular compartment. These observations support a model in which proliferative hyaloid cells migrate outside the vessel wall. This could serve as a primary driver of hyaloid vessels involution during the postnatal period rather than apoptotic clearance previously described.

**Fig. 1.**
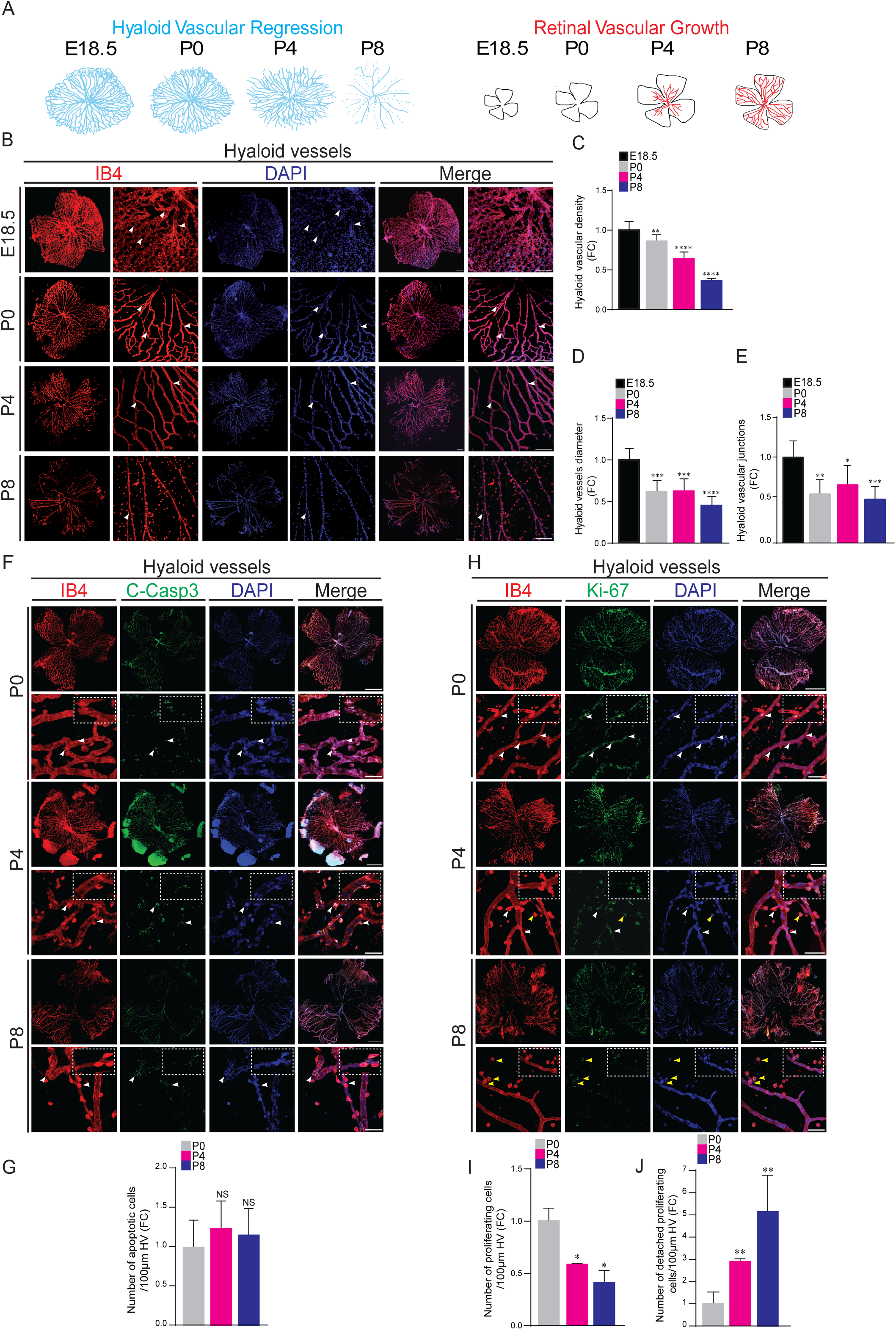
Cells relocation, not apoptosis, drives hyaloid pruning. (A) Schematic diagram of mouse hyaloid vessels regression and retinal vascular outgrowth at embryonic day 18.5 (E18.5) and postnatal days 0 (P0), 4 (P4), and 8 (P8). (B) Representative Isolectin B4 (IB4) and DAPI staining of E18.5, P0, P4 and P8 hyaloid vessels flatmounts show progressive loss of vascular density and branching. Arrowheads indicate vascular junctions. (C-E) Quantification of hyaloid vascular density, vessel diameter and vascular junctions (*n* = 6). Results are expressed as fold change (FC) normalized to E18.5 stage. (F) Representative confocal micrographs of P0, P4 and P8 hyaloid vessels flatmounts labeled with cleaved caspase 3 (C-Casp3), IB4 and DAPI reveal rare apoptotic cells with no significant temporal change, quantified per 100 µm of hyaloid vessel (HV) length (G) (*n* = 3). (H) Representative images of Ki-67, IB4 and DAPI staining of P0, P4 and P8 hyaloid vessels flatmounts show proliferating cells within (white arrowheads) and detached (yellow arrowheads) the vascular compartment, quantified per 100 µm of HV length (I, J) (*n* = 3). Results are expressed as FC normalized to P0 stage (G, I, J). Data are means ± SEM. Represented p-values are *≤0.05, **≤0.01, ***≤0.001, ****≤0.0001 from unpaired T-test. Non-significant (NS). Scale bars, 100 µm and 200 µm [for higher-magnification images in (B, F, H)].

### Cell-type specific contribution to postnatal hyaloid vessels regression

Given postnatal hyaloid vessels involution is characterized by sustained apoptotic activity and reduced proliferative capacity, we next examined the dynamics of hyalocytes, pericytes and endothelial cells to delineate their cellular contributions to vascular regression. We first examined hyalocytes, which have been implicated in apoptotic clearance and angiogenic regulation [8]. Immunostaining for the hyalocyte marker ionized calcium-binding adaptor molecule 1 (IBA1) revealed that the proportion of hyalocytes, whether vessel-associated or detached, remained unchanged between P0 and P8 (Fig. 2A, B). This distribution was accompanied by morphological heterogeneity. This distribution was accompanied by two consistent morphologies observed: elongated, vessel-adherent cells and smaller, rounded, detached cells (Fig. 2C, Fig. S1A). We next assessed pericyte dynamics, given their essential role in vessels stabilization and remodeling. Nerve/Glial Antigen 2 (NG2) immunostaining showed that pericyte numbers remained stable from P0 to P8 (Fig. 2D, E). At early stages (P0, P4), pericytes were predominantly vessel-associated, but by P8 a substantial fraction detaches from the blood vessel wall and appear in the vitreous (yellow arrowheads), consistent with stage-dependent loss of pericyte coverage that likely contributes to vessels destabilization (Fig. 2F). To complement NG2 staining, we assessed also α-smooth muscle actin (α-SMA), a marker of mature pericytes and vascular smooth muscle cells, which showed consistent expression in hyaloid vessels from P0 to P8 (Fig. S1D, E). This α-SMA staining is in accordance with the reported arterial identity of hyaloid vessels in mice [39]. Unlike the P8 retina which displays a specific α-SMA staining in the arteries but not veins (Fig. S1G) [40]. To determine whether this detachment pattern affect endothelial cells as reported before [41], we performed immunostaining for ETS-related gene (ERG1), a nuclear transcription factor specific to the endothelial lineage. Interestingly, while the total number of endothelial cells remained relatively constant from P0 to P8, their intravascular distribution became scattered (Fig. 2G, H). Similar to pericytes, ERG^positive^ cells transitioned from a vessel-adherent localization lining the vessel wall at early stages to extravascular pattern by P8 (Fig. 2G, I), implicating endothelial detachment as a cellular mechanism of regression. Collectively, these findings delineate a coordinated cellular remodeling process during hyaloid vessels regression. Rather than being eliminated through apoptosis, both pericytes and endothelial cells undergo progressive detachment and displacement from the vessel wall.

**Fig. 2.**
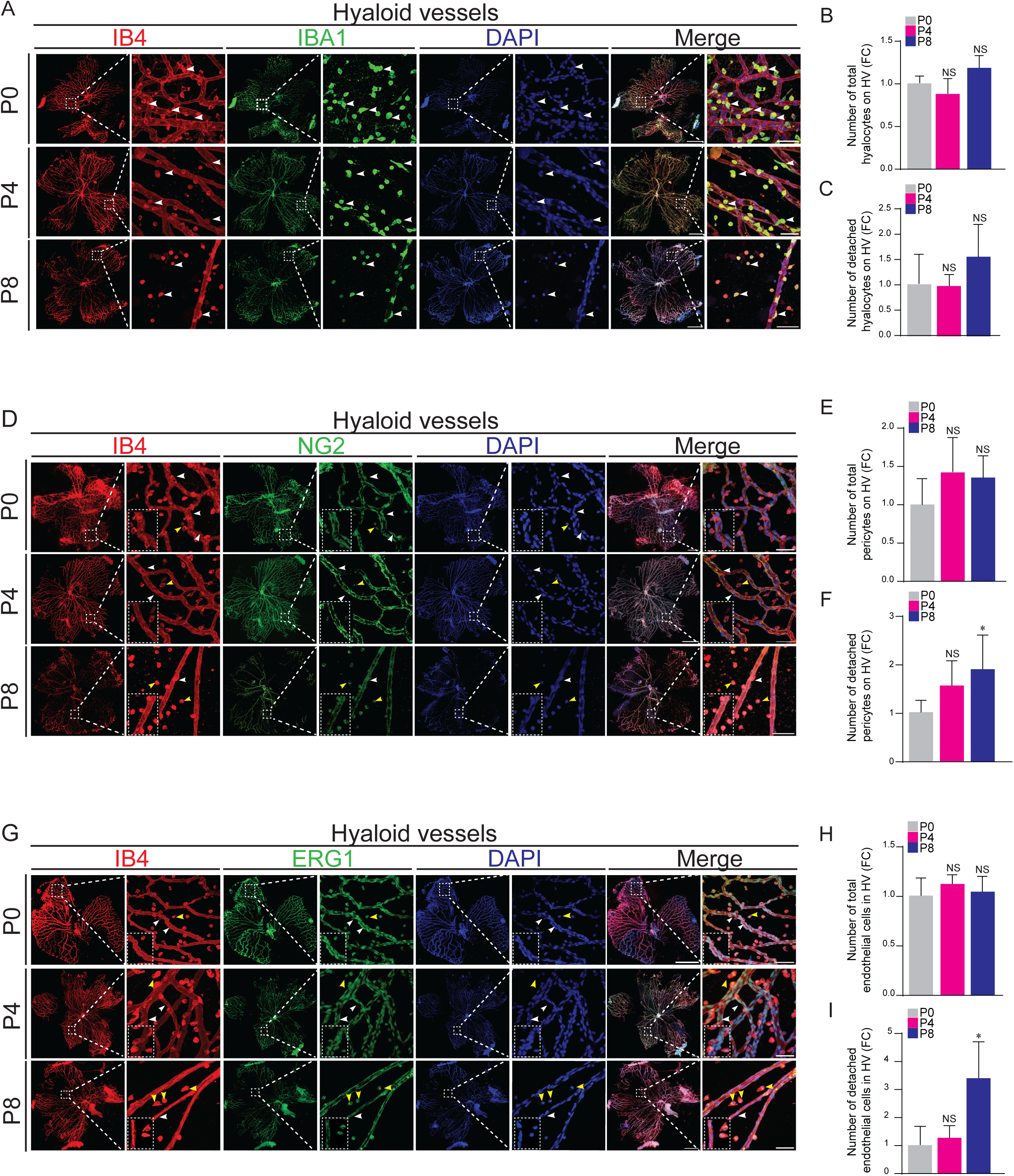
Cell-type specific contribution to postnatal hyaloid vessel regression. (A) Representative confocal micrographs of P0, P4 and P8 hyaloid vessels (HV) flatmounts stained with IBA1 (hyalocyte), IB4 and DAPI. White arrowheads indicate hyalocytes. (B) Quantification of total and (C) detached IBA1^positive^ hyalocytes, per 100 µm of HV length (*n* = 3). (D) Representative confocal immunofluorescence staining of NG2 (pericyte), IB4 and DAPI of P0, P4, and P8 hyaloid vessels flatmounts show morphological distinction between vessel-attached (white arrowheads) and detached pericytes (yellow arrowheads). (E) Quantification of the total and (F) detached NG2^positive^ pericytes per 100 µm of HV length (*n* = 3). (G) Representative P0, P4 and P8 hyaloid vessels flatmounts labeled with ERG1 (endothelial), IB4 and DAPI reveal a shift from adherent to dispersed patterns. Yellow arrowheads denote detached endothelial cells and white indicate attached endothelial cells. (H) Quantification of the total and (I) the detached ERG1^positive^ cells in 100 µm of HV length (*n* = 3). Results are expressed as fold change (FC) normalized to P0. Data are means ± SEM. Represented p-values are *≤0.05 from unpaired T-test. Non-significant (NS). Scale bars, 100 µm and 200 µm [for higher-magnification images in (A, D, G)].

### Hyaloid vascular regression involves endothelial-to-mesenchymal transition rather than endothelial apoptosis

To elucidate the cellular mechanisms underlying hyaloid vessels regression, we assessed the respective roles of endothelial apoptosis, inflammatory signaling, and fate transitions during vascular involution. Flow cytometric profiling of dissociated hyaloid vasculature, immunolabeled with PECAM1 (CD31; endothelial cell marker) and Annexin V (apoptosis), revealed that the relative frequencies of apoptotic endothelial (Annexin V⁺ CD31⁺) populations remained largely unchanged between P0 and P8 (Fig. 3A-C). Apoptotic profiles of non-endothelial populations (CD31⁻), including hyalocytes and pericytes, also remained unchanged across developmental stages (Fig. 3A, C). Consistent with this, the increase in detached extravascular endothelial and pericyte populations at P8 suggests that these cells migrate or delaminate from regressing vessels. Endothelial detachment in absence of increased cell death aligns with vascular remodeling mechanisms described in embryogenesis, where cells exit degenerating structures to maintain viability [42, 43].

**Fig. 3.**
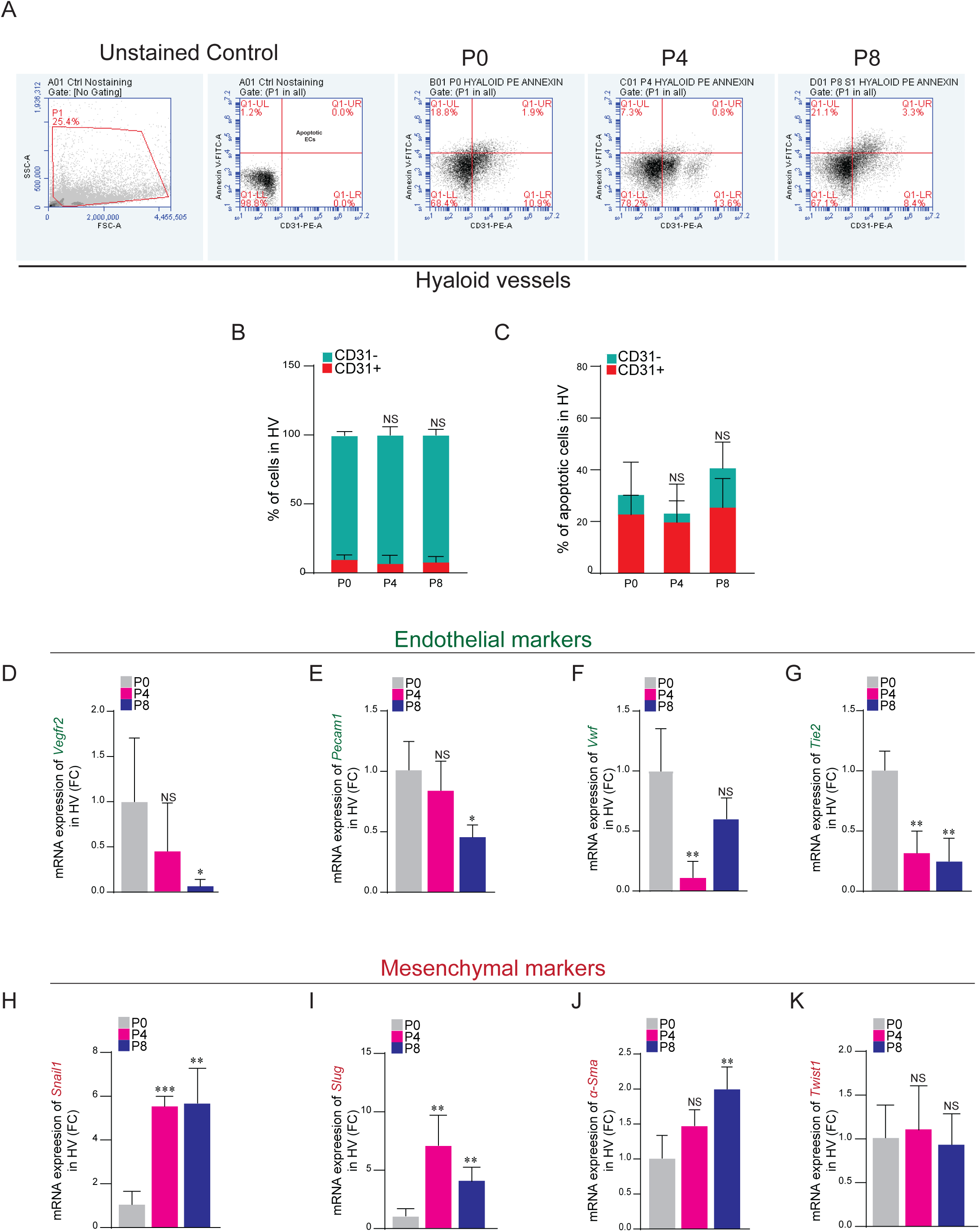
Hyaloid vascular regression involves endothelial-to-mesenchymal transition rather than endothelial apoptosis. (A) Flow cytometry analysis of apoptosis of endothelial cells (ECs) and non-ECs from hyaloid vasculature at P0, P4 and P8. Representative FACS plots show the gating strategy followed. Numerals shown in the plots indicate the percentages of the indicated cell population within each plot. (B) Quantification of total and (C) apoptotic (AnnexinV^+^) ECs (CD31⁺) and non-ECs (CD31⁻) cells populations. (D-G) Quantitative reverse transcription polymerase chain reaction (qRT-PCR) shows downregulation of endothelial identity markers *Vegfr2*, *Pecam1*, *Vwf* and *Tie-2* during hyaloid regression. (H-K) qRT-PCR analysis shows induction of EndoMT-associated transcription factors: *Snail1*, *Slug* and *α-Sma* in hyaloid from P0 to P8. *Twist* expression remains unchanged (K). β-actin was used as a reference gene. Results are expressed as fold change (FC) normalized to P0 stage (D-K). Data are means ± SEM. Represented p-values are *≤0.05, **≤0.01, ***≤0.001 from unpaired T-test. *n* = 3-6 for *Vegfr2*; *n* = 3-4 for *Pecam1, Twist* and *Snail*; *n* = 3-5 for *Vwf* and *Tie2*; *n* = 4 for *Slug* and *α-Sma* (E-L). Non-significant (NS).

In line with this, we observed a rise in detached endothelial cells without a concomitant increase in apoptotic activity. We therefore investigated whether endothelial cells undergo endothelial-to-mesenchymal transition (EndoMT), a process that facilitates migration through loss of endothelial identity and acquisition of mesenchymal characteristics. Quantitative reverse transcription polymerase chain reaction (qRT-PCR) analyses revealed a marked downregulation of endothelial markers *Vegfr2*, *Pecam1*, and *Tie2* during hyaloid regression from P0 to P8. *Vwf* was significantly reduced at P4 and showed a further, though non-significant, decline by P8 (Fig. 3D-G). Loss of endothelial markers coincided with strong induction of mesenchymal factors *Snail1*, *Slug* and *α-Sma*, while *Twist*1 remained unchanged (Fig. 3H-K). This indicates selective engagement of the EndoMT program, consistent with models where *Snail1* and *Slug* drive an early, plastic transition that can revert, whereas *Twist1* typically enforces a sustained, pathological mesenchymal state [44–48]. We next profiled hyalocyte inflammatory states during hyaloid regression. Cytokine analysis showed significant upregulation of *Il-1β* at P8 and *Tnf-α* at P4, consistent with a pro-inflammatory M1 phenotype (Fig. S2A, B). By contrast, the M2 markers transcripts were not detected at any stage. These findings suggest that alternative anti-inflammatory or M2-like programs are not engaged during hyaloid regression, pointing instead to a predominant M1-polarized state that may drive vessel destabilization and regression.

Together, these findings indicate that hyaloid vascular regression is mediated by endothelial reprogramming and inflammatory activation, in addition to apoptosis. Endothelial cells detach from regressing vessels, likely via EndoMT, while hyalocytes sustain a pro-inflammatory environment that contributes to hyaloid vessels regression and eye tissue remodeling.

### Notch1 pathway expression in hyaloid regression and proliferation-migration control in vitro

Prompted by our observations of EndoMT and endothelial-like delamination during hyaloid vessels regression, we set out to investigate upstream regulatory pathways that may govern these dynamic cellular transitions. Notch1, known for its roles in vascular stabilization and EndoMT [49–51], was next investigated for a previously unrecognized function in hyaloid vessel regression. To address this, we first assessed the temporal expression of Notch1, its ligands, and canonical downstream targets in hyaloid vessels during the regression window (P0-P8).

Western blot analysis of isolated hyaloid vessels showed a progressive increase in total Notch1 protein level from P0 to P8, peaking at P4, with the cleaved active form (Notch1 intracellular domain, N1ICD) also maximal at P4 (Fig. 4A). DLL4 and JAG1 displayed similar temporal dynamics at the protein level, with expression transiently peaking at P4 before returning to baseline by P8 (Fig. 4aA). Consistent with these dynamics, qRT-PCR analysis revealed a transient increase in Notch1 mRNA at P4, which declined to baseline by P8 (Fig. 4B). By contrast, *Dll4* transcript levels also decreased by P8 (Fig. 4C), whereas Jag1 expression significantly increased at this stage (Fig. 4D), highlighting divergent ligand regulation during hyaloid regression. Transcriptional profiling of canonical Notch targets revealed sustained upregulation of *Hes1* at both P4 and P8 (Fig. 4E), consistent with ongoing Notch1 activity. In contrast, expression of *Hey2* revealed upregulation at P4 and downregulation at P8, while *Hes5* was significantly downregulated at these time points (Fig. S3A, B), suggesting distinct, possibly non-redundant roles of Notch target genes during this process.

**Fig. 4.**
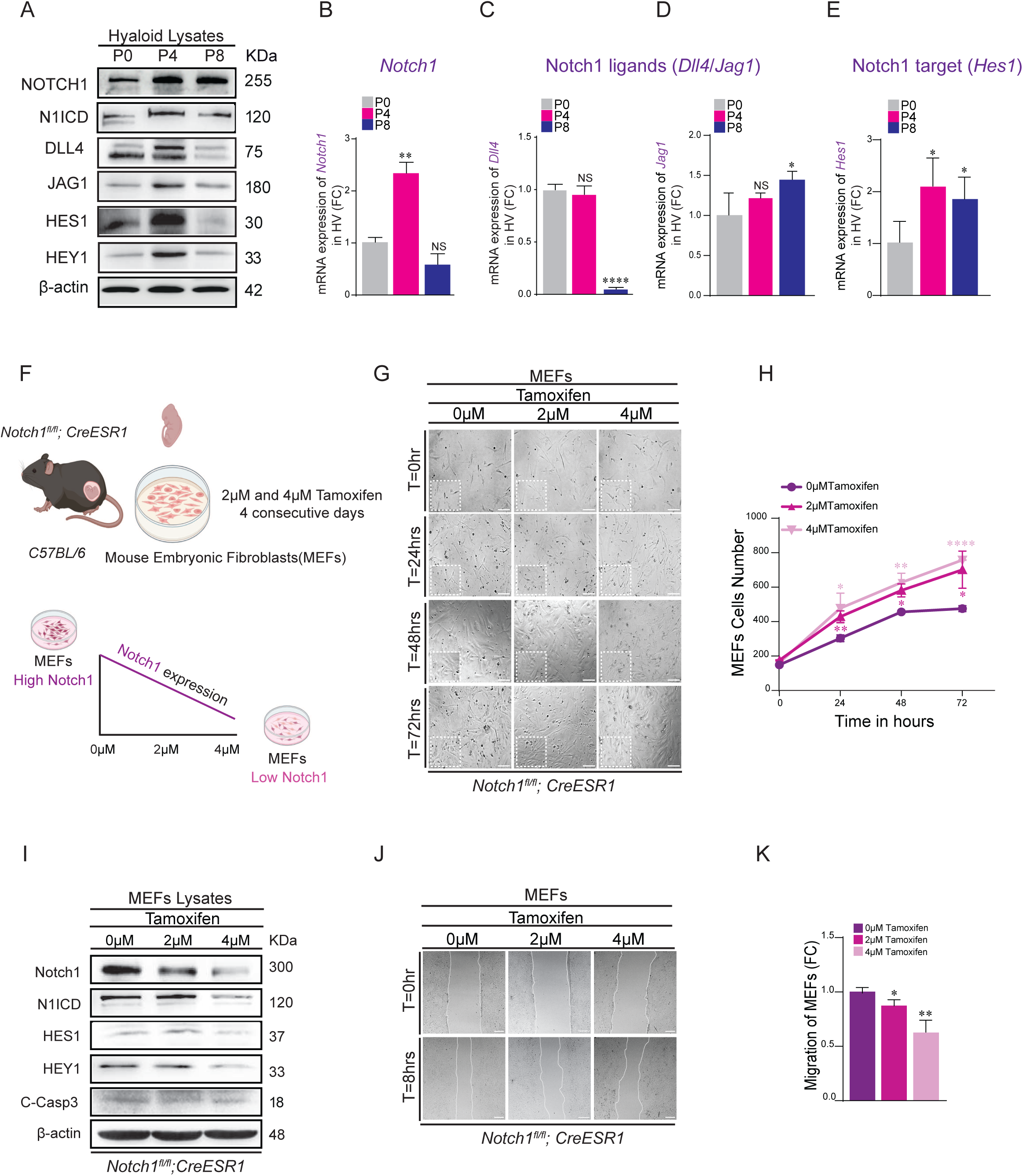
Notch1 pathway expression in hyaloid regression and proliferation-migration control in vitro. (A) Western blot of hyaloid cell lysates from P0, P4 and P8 shows the protein expression of Notch1, its active form (Notch1 intracellular domain, N1ICD), its ligands DLL4 and JAG-1 and its target genes HES1 and HEY1. β-actin was used as a loading control. (B) qRT-PCR analysis shows the levels of *Notch1*, (C) *Dll4*, (D) *Jag1* and (E) *Hes1* transcripts in hyaloid vessels (HV) at P0, P4, and P8. β-actin was used as a reference gene. Results are expressed as fold change (FC) normalized to P0 stage (F) Schematic depiction of tamoxifen-inducible *Notch1* knockout mouse model *Notch1^fl/fl^; CreESR1* in which *Notch1* is deleted in mouse embryonic fibroblasts (MEFs) in a dose-dependent manner (2 and 4 µM tamoxifen). (G) Representative images of *Notch1^fl/fl^; CreESR1* MEFs from the time-course proliferation assay at the initial time-point (0 hours) and after 24, 48 and 72 hours, treated and non-treated with 2 and 4 µM tamoxifen, show increased growth in a dose-dependent manner, quantified in (H). (I) Western blot of *Notch1^fl/fl^; CreESR1* MEFs lysates treated or non-treated with 2 and 4 µM tamoxifen shows decrease of Notch1, N1ICD, HES1 and HEY1 in a dose-dependent manner and constant protein expression of apoptotic marker C-Casp3. β-actin was used as a loading control. (J) Representative images from the wound edge at the initiation of imaging (0 hours) and 8 hours after wounding of Notch1 knockout MEFs monolayers, treated with 2 and 4 µM tamoxifen show a dose-dependent decrease in migration compared to non-tamoxifen treated, Cre positive MEFs control, quantified in (K). Data are means ± SEM. Represented p-values are *≤0.05, **≤0.01, ****≤0.0001 from one-way ANOVA with unpaired T-test. *n* = 3-5 for *Notch1*; *n* = 3-7 for *Dll4*; *n* = 5-8 for *Jag1*; *n* = 3-4 for *Hes1* (B-E)*. n* = 3 biologically independent experiments of proliferation and migration (G-K). Non-significant (NS). Scale bars, 200 µm (G) ; 100 µm (J).

To assess the broad role of Notch1 in regulating proliferation, migration and apoptosis, we examined both fibroblasts, which provide a tractable model for studying stromal-like contributions to remodelling processes (e.g. during wound healing and fibrosis) [52] and endothelial cells, which are central to vascular regression through processes such as EndoMT [41, 53]. For fibroblasts, we used mouse embryonic fibroblasts (MEFs) derived from a tamoxifen-inducible Notch1 knockout mouse model (*Notch1^fl/fl^*; *CreESR1*), enabling precise temporal control of Notch1 deletion *in vitro* (Fig. 4F). Interestingly, Notch1 loss doubled MEF proliferation over 72 hours following treatment with 2 or 4 µM tamoxifen compared with untreated controls (Fig. 4G, H). This proliferative response correlated with a dose-dependent reduction in Notch1 protein and diminished expression of downstream targets HES1 and HEY1, as shown by western blotting (Fig. 4I). Furthermore, immunoblot analysis confirms that Notch1 knockdown did not trigger apoptosis, as C-Casp3 remained at low level suggesting that Notch1 silencing does not compromise cell viability under these experimental conditions (Fig. 4I).

In contrast to its suppressive effect on proliferation, Notch1 was required for MEF migration. Scratch-wound assays showed a dose-dependent defect in wound closure after tamoxifen-mediated Notch1 ablation, with significantly reduced migration at both 2 and 4 µM compared with untreated cells. (Fig. 4J, K). To corroborate these findings, we pharmacologically inhibited Notch signalling using the γ-secretase inhibitor DAPT, which prevents N1ICD release. Similar to the genetic knockout, DAPT treatment markedly impaired MEF migration (∼2-fold reduction) while strongly enhancing proliferation (∼6-fold increase) over 72 hours (Fig. S3D, E). Importantly, these effects occurred without changes in C-Casp3, confirming a Notch1-specific mechanism independent of apoptosis (Fig. S3H).

Consistent with MEFs, pharmacological inhibition of Notch signalling in bovine aortic endothelial cells (BAECs) significantly impaired migration (∼1.6-fold reduction) and suppressed key pathway effectors, including N1ICD and HES1, without evidence of apoptosis. (Fig. S3I-K). Notch1 signalling thus integrates proliferative restraint with migratory competence, positioning it as a central regulator of mesenchymal-endothelial behaviour and tissue remodelling during hyaloid vessel regression. Together, these findings identify Notch1 signalling integrates proliferative restraint with migratory competence, establishing it as a central regulator of endothelial mesenchymal behaviour in vitro and a potential driver of tissue remodelling during hyaloid vessel regression.

### Endothelial-specific Notch1 deletion *in vivo* disrupts hyaloid vascular regression and leads to persistence

To investigate the functional role of Notch1 *in vivo* during hyaloid vessels regression, we used a tamoxifen-inducible, endothelial cell-specific Notch1 knockout mouse model *(Notch1^fl/fl^; Cdh5-CreERT2*) identified as *Notch1^cKO^*. This mouse model allows for temporally controlled deletion of Notch1 specifically in vascular endothelial cells upon administration of tamoxifen. To target the period of active hyaloid vessels remodeling, tamoxifen was administered via intragastric injection to neonatal pups from P1 to P3, and hyaloid vessels were collected at P8 as demonstrated in the model (Fig. 5A). Efficient *Notch1* deletion was confirmed in hyaloid vessels, where mRNA levels were reduced to near-background in *Notch1^cKO^* hyaloids relative to tamoxifen-treated, Cre-negative littermates (*Notch1^cWT^)* control P8 (Fig. 5B). This model provided a robust platform to dissect the endothelial-specific function of Notch1 in postnatal vascular regression without confounding developmental lethality. In stark contrast to the regressed network observed in P8 *Notch1^cWT^* control, immunofluorescence staining of flatmounted P8 hyaloid vessels with IB4 revealed robust persistence in *Notch1^cKO^* littermates (Fig. 5C). Quantitative morphometric analysis showed a 2.5-fold increase in vascular density, a 2-fold increase in vessels diameter and a 2.5-fold increase in branching *Notch1^cKO^* hyaloids compared to controls (Fig. 5D-F), indicating that Notch1 deletion disrupts normal hyaloid vessels regression and stabilization.

**Fig. 5.**
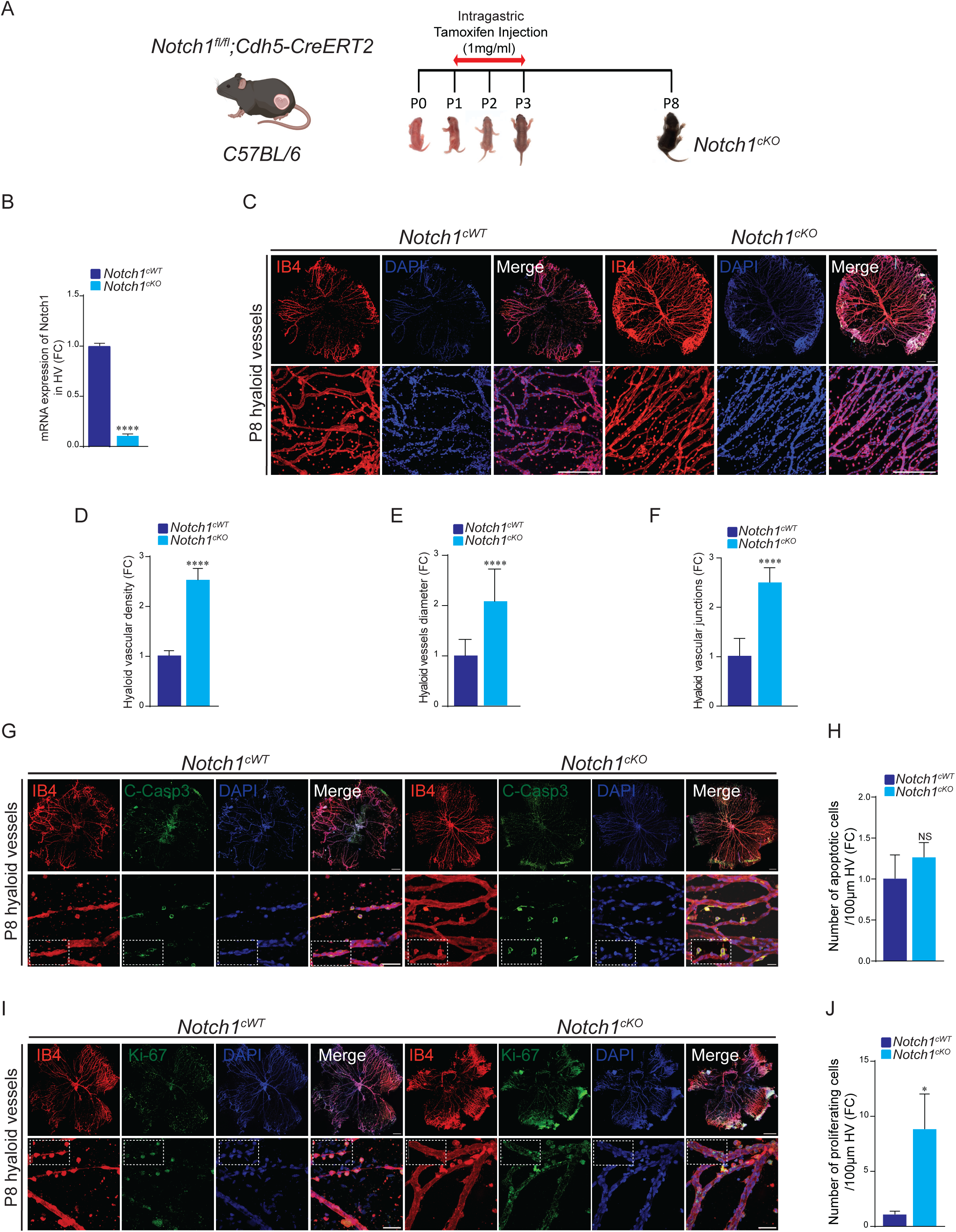
Endothelial-specific Notch1 deletion *in vivo* disrupts hyaloid vascular regression and leads to persistence. (A) Schematic diagram of tamoxifen-induced Notch1 deletion in ECs in *Notch1^fl/fl^; Cdh5-CreERT2 C57BL/6* transgenic mice. (B) Validation of *Notch1* deletion by qRT-PCR in *Notch1^cKO^*versus *Notch1^cWT^* hyaloid vessels (HV). β-actin was used as a reference gene. (C) Representative confocal micrographs of P8 hyaloid vessels flatmounts in *Notch1^cKO^* and *Notch1^cWT^* mice immunostained with IB4 and DAPI. (D) Quantification of hyaloid vascular density, (E) vessels diameter, and (F) number of junctions in *Notch1^cKO^*versus *Notch1^cWT^* (*n* = 6). (G) Representative confocal micrographs of P8 hyaloid vessels flatmounts in *Notch1^cKO^* versus *Notch1^cWT^* labeled with cleaved caspase 3 (C-Casp3), IB4 and DAPI reveal rare apoptotic cells with no significant change, quantified in (H) per 100µm hyaloid vessels length (*n* = 3). (I) Representative images of Ki-67, IB4 and DAPI staining of P8 hyaloid vessels flatmounts show higher proliferating cells in *Notch1^cKO^* versus *Notch1^cWT^*, quantified in (J) (*n* = 3). Data are means ± SEM. Represented p-values are *≤ 0.05, ****≤0.0001 from unpaired T-test. *n* = 3-4 for *Notch1* (B). Non-significant (NS). Scale bars, 100µm and 200µm [for higher-magnification images in (C, G, I)].

To determine whether altered cell turnover contributed to this phenotype, we examined apoptosis and proliferation of hyaloid vessels in *Notch1^cKO^*. Endothelial-specific deletion of Notch1 did not affect basal apoptosis, as C-Casp3 staining revealed similar numbers of apoptotic cells between *Notch1^cKO^* and *Notch1^cWT^* vessels at P8 (Fig. 5G-H), suggesting that the persistent vasculature was not due to impaired apoptosis. However, Ki-67 staining revealed a striking 9-fold increase in proliferating cells within persistent hyaloid vessels of *Notch1^cKO^* mice compared with the sparse, largely extravascular proliferating cells in *Notch1^cWT^*vessels at P8 (Fig. 5I, J). EdU incorporation further confirmed this phenotype, showing 4-fold increase in DNA-synthesizing cells in *Notch1^cKO^* vessels (Fig. S4A, B). Together, these data establish that Notch1 constrains cell proliferation during hyaloid regression, and that loss of this restraint underlies the persistence phenotype. Moreover, endothelial-specific deletion of Notch1 impaired retinal vascular development, as IB4 staining of flatmounted P8 retinas revealed a marked delay in superficial vascular layer formation compared with *Notch1^cWT^* controls (Fig. S4C). Quantitative morphometric analysis showed a 6-fold decrease in vascular area, and 5-fold decrease in branching in *Notch1^cKO^* retinal vasculature compared with controls (Fig. S4D, E). Furthermore, 3D reconstruction of IB4-stained retinas showed that *Notch1^cKO^* mice failed to fully develop the deep vascular plexus at P14 and P21, in contrast to *Notch1^cWT^* controls (Fig. S4F, G) [54]. Together, these results confirm that Notch1 is essential for hyaloid vessels involution *in vivo*. Its endothelial-specific deletion results in excessive proliferation and stabilization of the hyaloid network, providing direct evidence that Notch1 signaling is a key driver of physiological vascular regression during ocular development.

### Notch1 deficiency prevents endothelial like delamination and represses EndoMT during hyaloid vascular persistence

To investigate the cellular basis of hyaloid vessels persistence in *Notch1^cKO^* mice, we examined endothelial behavior at P8, a developmental stage when *Notch1^cWT^* hyaloid network normally regress, exhibiting nearly 70% reduction in vessels density compared to E18.5 (Fig. 1). We performed flatmounted immunostaining using ERG1, IB4, and DAPI to compare endothelial distribution between *Notch1^cKO^* and *Notch1^cWT^* hyaloid vasculature (Fig. 6A). *Notch1^cKO^* flatmounted hyaloids showed confined ERG^positive^ cells within the IB4 stained blood vessels, with almost complete absence of extravascular endothelial cells compared to the *Notch1^cWT^* control (Fig. 6A). Quantification revealed a significant elevation in endothelial cell number in *Notch1^cKO^* hyaloids compared to *Notch1^cWT^* (Fig. 6B), while detached (non-vessel-associated) ERG^positive^ cells were completely absent in the knockout (Fig. 6C). These findings suggest that Notch1 is required for endothelial detachment and redistribution to the extravascular areas during postnatal hyaloid vessels regression. Given our earlier findings that hyaloid regression is accompanied by EndoMT, we next assessed whether Notch1 is required for this process. mRNA profiling of P8 hyaloid vessels showed that *Notch1^cKO^* samples upregulated endothelial markers (*Vegfr2*, *Pecam1*, *Vwf*, and *Tie2;* Fig. 6D-G), consistent with endothelial stabilization, while mesenchymal markers (*Snail1*, *Slug*, *α-Sma* and *Twist1;* Fig. 6H-K) were significantly reduced compared with *Notch1^cWT^*. These transcriptional changes indicate that Notch1 is required to activate EndoMT-associated genes during regression, and that its loss locks endothelial cells in a stable, vessel-adherent state, thereby promoting vascular persistence. These transcriptional changes indicate that Notch1 activity is required for the induction of EndoMT-associated genes during physiological regression. Collectively, these data demonstrate that Notch1 deficiency locks endothelial cells in a vessel-adherent, identity-stable state, preventing their redistribution and contributing to persistent vascular structures. This uncovers a mechanistic link between Notch1 signaling, EndoMT, and endothelial repositioning during developmental vessels involution. To determine whether Notch1 deficiency also impacts supporting mural cells, we examined pericyte distribution in P8 hyaloid vessels. Flat-mounted hyaloids from *Notch1^cKO^*and *Notch1^cWT^* were immunostained for NG2, IB4, and DAPI (Fig. S5A). Pericyte number was significantly elevated in *Notch1^cKO^* hyaloids (Fig. S5B). Unlike controls, where many pericytes delaminated into the extravascular space, most *Notch1^cKO^* pericytes remained vessel-associated (Fig. S5C, yellow arrowheads), indicating defective migratory behavior.

**Fig. 6.**
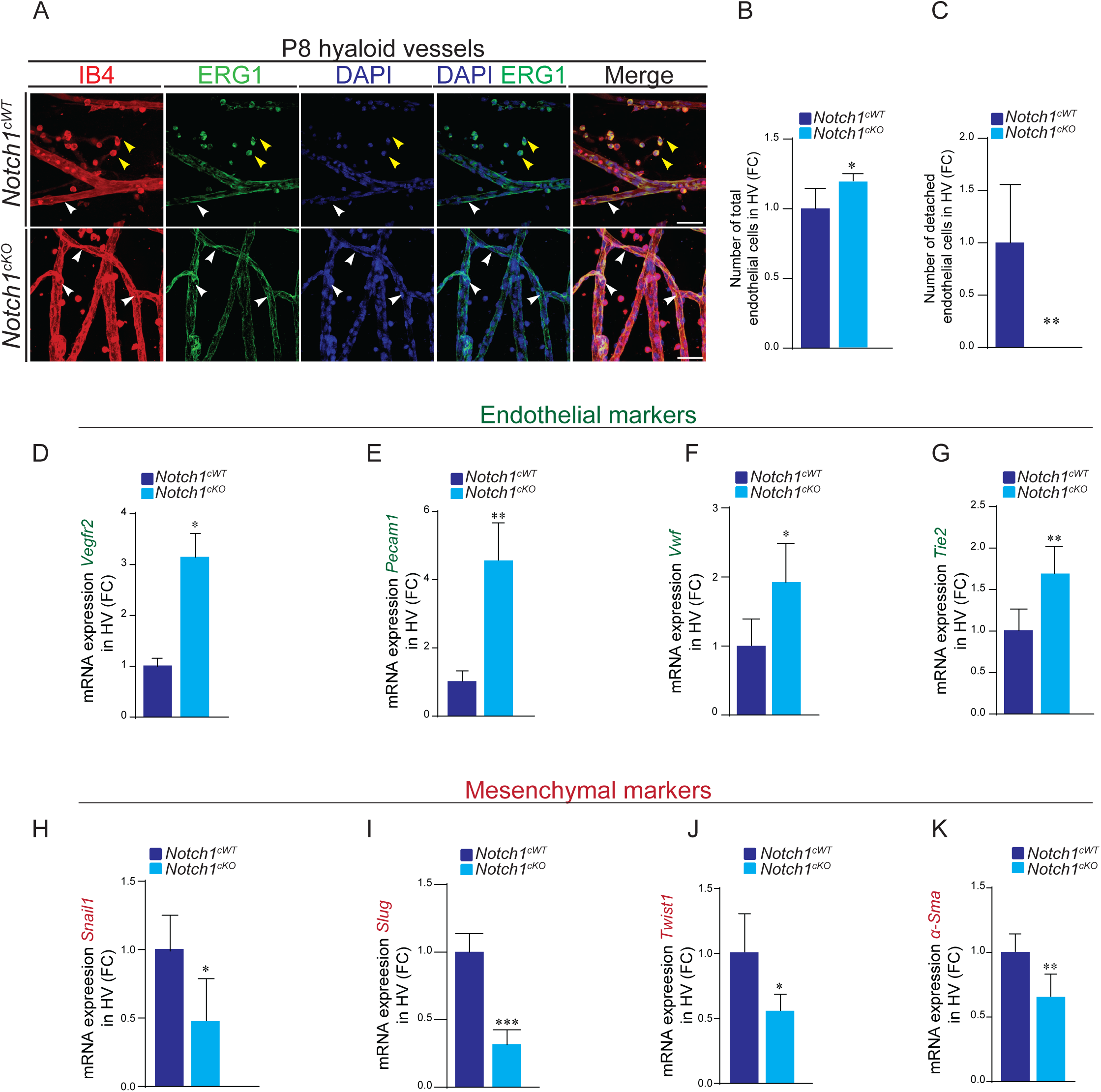
Notch1 deficiency prevents endothelial like delamination and represses EndoMT during hyaloid vascular persistence. (A) Representative confocal micrographs of P8 hyaloid vessels (HV) flatmounts in *Notch1*^cKO^ versus *Notch1^cWT^* immunostained with ERG1, IB4 and DAPI. Yellow arrowheads indicate detached endothelial cells, and white arrowheads indicate attached endothelial cells (*n* = 3). (B) Quantification of total and (C) detached ERG1^positive^ cells in *Notch1*^cKO^ compared to *Notch1^cWT^* hyaloid vessels (*n* = 3). (D) qRT-PCR shows upregulation of endothelial identity markers *Vegfr2*, (E) *Pecam1*, (F) *Vwf* and (G) *Tie2* in *Notch1^cKO^* versus *Notch1^cWT^*. (H) qRT-PCR analysis shows reduction of EndoMT-associated transcription factors *Snail1*, (I) *Slug*, (J) *Twist* and (K) α-*Sma* in *Notch1^cKO^* hyaloid vessels versus *Notch1^cWT^*. β-actin was used as a reference gene. Results are presented as fold change (FC) relative to P8 *Notch1^cWT^*. Data are means ± SEM. Represented p-values are *≤ 0.05, **≤ 0.01, ***≤ 0.001 from unpaired T-test. n = 3 for *Vegfr2*; n = 3-4 for *Pecam1*; n = 4 for *Vwf*; n = 5 for *Tie2;* n = 4-6 for *Snail1*; n = 3-4 for *Slug*; n = 4-5 for *Twist1*; n = 4 for *α-Sma*. Scale bars, 200µm.

To assess if inflammatory signaling is altered in persistent hyaloid vasculature, we analyzed hyalocyte dynamics using IBA1 immunostaining. Both total IBA1^positive^ hyalocyte number (Fig. S5E), and detached hyalocytes were reduced by more than half in *Notch1^cKO^* P8 hyaloids compared with controls (Fig. S5D-F). This phenotype resembles the early postnatal stage (P0), when hyalocytes remain predominantly vessel-associated, suggesting that persistent vasculature in the absence of Notch1 retains immature, pre-regression characteristics. (Fig. S1C). To explore macrophage activation states, we quantified the mRNA levels of key M1 and M2 inflammatory markers. *Il-1β* and *Tnf-α*, gene expression remained unchanged in *Notch1^cKO^* hyaloid vessels (Fig. S5G, H), indicating a sustained pro-inflammatory microenvironment. qRT-PCR failed to detect M2-associated markers in *Notch1^cKO^* hyaloids demonstrating that anti-inflammatory resolution programs are not activated under these conditions. Together, loss of Notch1 prevents EndoMT and perivascular cell redistribution while maintaining a pro-inflammatory milieu, thereby stabilizing endothelial identity and driving persistent hyaloid vasculature.

### Notch1-Wnt transcriptional crosstalk orchestrates endothelial plasticity during hyaloid vessel regression

Given the established role of the Wnt receptors LRP5, LRP6, and FZD4 in hyaloid vessel regression, we investigated whether Notch1 loss impacts their expression [55]. qRT-PCR analyses revealed a marked (>2-fold) reduction in *Lrp5*, *Lrp6*, and *Fzd4* expression in *Notch1^cKO^* hyaloid vessels relative to littermate controls (Fig. 7A-C). To further probe the interplay between Wnt and Notch signaling, we re-analyzed publicly available single-cell RNA sequencing (scRNA-seq) datasets from *Frizzled 5* (*Fz5^-/-^*) mutants at P3, which display persistent hyaloid vasculature [56]. Strikingly, loss of *Fz5* was accompanied by a significant suppression of Notch1-associated pathways, reinforcing a reciprocal requirement of Wnt and Notch signaling in hyaloid vessel regression (Fig. 7D-E).

**Fig. 7.**
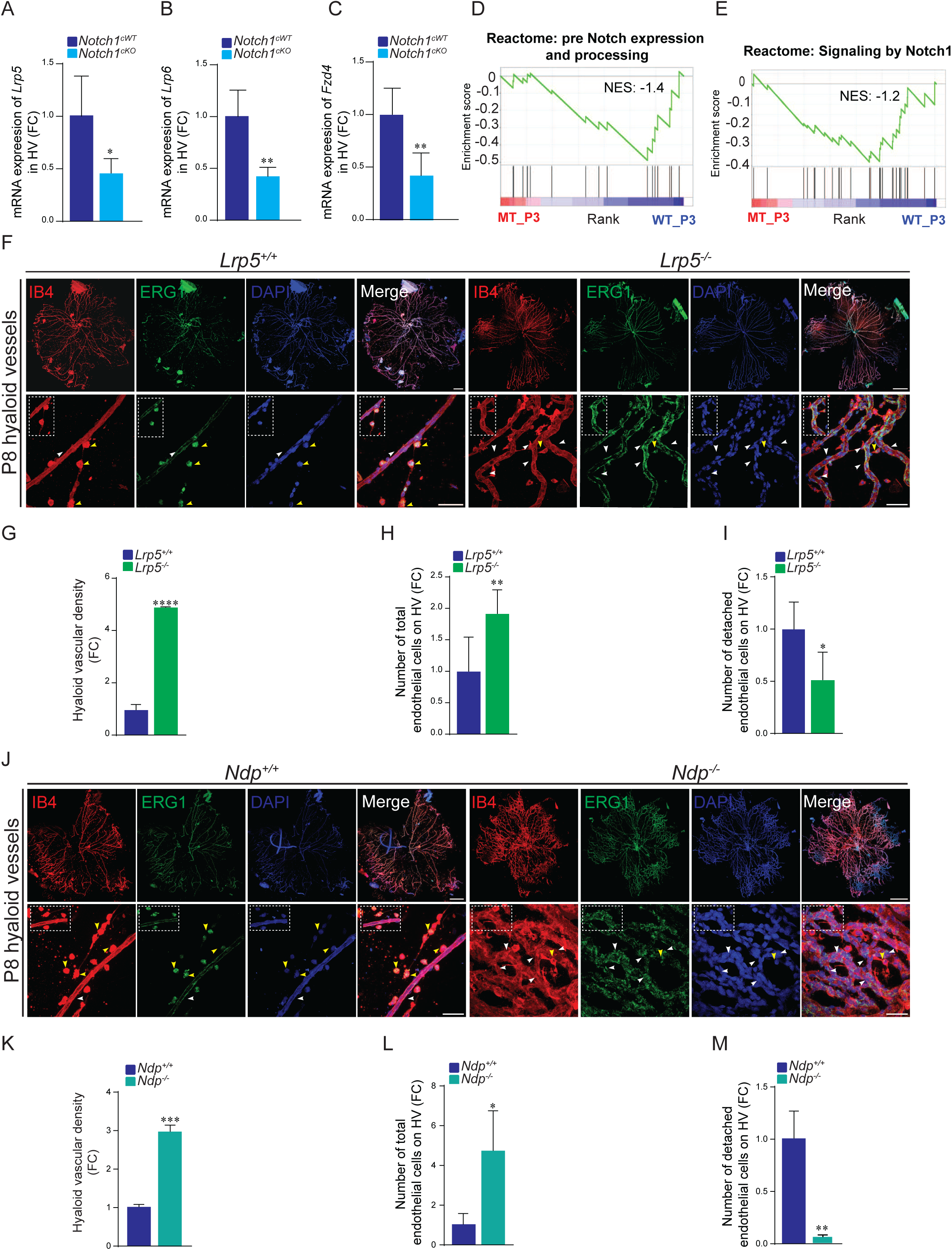
Notch1-Wnt transcriptional crosstalk orchestrates endothelial plasticity during hyaloid vessel regression. (A) qRT-PCR analysis shows transcripts expression of *Lrp5*, (B) *Lrp6*, and (C) *Fzd4* in P8 *Notch1^cKO^* versus *Notch1^cWT^* hyaloid vessels. β-actin was used as a reference gene. (D, E) GSEA bar plots displaying normalized enrichment scores (NES) from Reactome pathway analysis (FDR q < 0.05) of endothelial cell scRNA-seq data. In postnatal day 3 (P3) *Fz5^-/-^* mutants, pathways associated with pre-Notch expression, processing, and Notch1 signaling are downregulated relative to *Fz5^+/+^* controls. (F) Representative confocal micrographs of P10 hyaloid vessels (HV) flatmounts from *Lrp5^-/-^* and *Lrp5^+/+^* mice immunostained with ERG1, IB4 and DAPI *(n* = 2-3). (G) Quantification of hyaloid vascular density, (H) total and (I) detached ERG1^positive^ endothelial cells in *Lrp5^-/-^* versus *Lrp5^+/+^* hyaloid vessels. (J) Confocal images of ERG1, IB4 and DAPI immunostaining in P8 *Ndp^-/-^* versus *Ndp^+/+^* hyaloid vessels (*n* = 2-3). White arrowheads indicate attached endothelial cells, and white arrowheads indicate attached endothelial cells. (K) Quantification of hyaloid vascular density, (L) total and (M) detached ERG1^positive^ endothelial cells in *Ndp^-/-^* versus *Ndp^+/+^* hyaloid vessels. Results are expressed as fold change (FC) relative to *Lrp5^+/+^*, *Ndp^+/+^* and *Notch1^cWT^*control hyaloid vessels. Data are means ± SEM. Represented p-values are *≤ 0.05, **≤ 0.01, ***≤ 0.001, ****≤0.0001 from unpaired T-test. Non-significant (NS). Scale bars, 100µm and 200µm [for higher-magnification images in (C, D)].

To investigate the implication EndoMT as a general mechanism during hyaloid regression, we performed hyaloid flatmounted of the Norrie knockout (*Ndp^-/-^*) and Lrp5 knockout (*Lrp5^-/-^*) hyaloids. We confirmed their persistent phenotype as already reported (Fig. 7F, J), confirmed with the quantification of hyaloid vascular density (Fig. 7G, K) [57, 58]. Furthermore, immunostaining for the endothelial marker ERG1 recapitulated the vascular phenotype observed in *Notch1^cKO^* hyaloids (Fig. 6A). Notably, both *Lrp5^-/-^* and *Ndp^-/-^* mutants exhibited a striking accumulation of ERG^positive^ nuclei restricted within IB4^positive^ vascular structures, with a near-complete loss of extravascular endothelial cells (Fig. 7F, J, yellow arrowheads). Quantification of total and detached ERG^positive^ cells in hyaloid vessels confirmed this endothelial defect (Fig. 7H, I, L, M). This phenotype points to a failure of endothelial cells to detach from the vessel core, highlighting a shared requirement of Wnt-Notch signaling components for endothelial delamination during hyaloid regression. While it remains unresolved whether this reflects direct regulatory cross-talk, these findings point to a potential transcriptional coupling between Notch1 and Wnt pathways in ocular vascular remodeling, with precise molecular interactions yet to be defined [59, 60].

## Discussion

Developmental regression of the hyaloid vasculature is a tightly choreographed process essential for ocular maturation, yet the molecular cues governing its involution remain poorly defined. Here, we identify Notch1 as a central regulator of hyaloid vessel regression, coordinating proliferative restraint, endothelial delamination, and signaling crosstalk with Wnt.

Our data reveal that Notch1 activity rises during the regression phase, marked by induction of DLL4, JAG1, and the downstream effector HES1. Genetic deletion of Notch1 disrupted regression and led to vascular persistence, characterized by excessive branching, vessel dilation, and increased endothelial density. Notably, these changes occurred independently of apoptosis, establishing Notch1 as a gatekeeper of proliferation rather than a driver of cell death during postnatal regression. This aligns with its broader role in angiogenesis, where Notch1 antagonizes VEGF-driven expansion while enforcing vascular quiescence [61].

This work identifies a previously unrecognized mechanism underlying hyaloid vessel regression. Notch1 limits endothelial proliferation while driving their detachment from the vascular wall. In the absence of Notch1, endothelial identity programs (*Vegfr2*, *Pecam1*, *Vwf*, *Tie2*) persist and mesenchymal markers (*Snail1*, *Slug*, *Twist1*, *α-SMA*), remain suppressed, reflecting an aborted EndoMT program. Such reversible hybrid states are increasingly recognized as critical for tissue remodeling, as seen in cardiac valve morphogenesis and pulmonary artery development [62–64]. In Notch1-deficient vessels, pericytes also failed to detach, and hyalocytes adopted a sustained pro-inflammatory state, suggesting that Notch1 coordinates multicellular remodeling events required for vascular involution.

A striking feature of *Notch1* loss was the downregulation of Wnt signaling components *Lrp5*, *Lrp6*, and *Fzd4*, receptors previously implicated in retinal angiogenesis and hyaloid regression [65–69]. Together with re-analysis of *Fz5* mutant scRNA-seq data, which revealed diminished Notch pathway activity, our results point to a bidirectional reinforcement between Wnt and Notch signaling during hyaloid regression. Although the precise transcriptional mechanisms remain to be defined, these findings raise the possibility that Notch1 directly sustains Wnt receptor expression, establishing a positive feedback loop essential for endothelial plasticity and vessel regression.

In sum, our study establishes Notch1 as a master regulator of hyaloid vessel regression, acting not through apoptosis but by enforcing proliferative arrest. It enables EndoMT and sustains Wnt-Notch crosstalk. This coordinated program culminates in a delamination-like phenotype essential for vascular involution. The failure of this process in Notch1-deficient and mice and the mouse models of PFV (*Ndp* and *Lrp* deficient mice) highlights how disruption of Notch-Wnt coupling may underlie persistent fetal vasculature and related ocular pathologies characterized by impaired vessel regression.

## Material and Methods

### Murine model

The manipulations carried out on mice as part of this project were approved by the Institutional Animal Protection Committee (CIPA) of the University of Quebec in Montreal (UQAM) accredited by the Canadian Council on Animal Care (CCAC). For inducible endothelial deletion of *Notch1, Notch1^fl/fl^* and *Cdh5-CreERT2* mice were crossed to obtain (*Notch1^fl/fl^; Cdh5-CreERT2*) mice.

To knock-out *Notch1* ubiquitously, *Notch1^fl/fl^* and *CreESR1* mice were crossed to generate (*Notch1^fl/fl^; CreESR1)* mice. These mice were used for mouse embryonic fibroblasts (MEFs) extraction. *Cre-ESR* and *Notch1^fl/fl^* mice were kindly provided by Przemyslaw Sapieha while *Cre-Cdh5* mice by Bruno Larrivée. Mice strains were of *C57BL/6J* genetic background. Genotyping was performed by polymerase chain reaction (PCR) analysis of genomic DNA extracted from ear punches. Mice with the correct genotype were selected for subsequent experiments.

C57BL/6J wild-type (*WT*), Lrp5 knock-out (*Lrp5^-/-^*) (stock # 005823) and Ndp knock-out (*Ndp^-/-^*) (stock # 012287) mice were from Jackson Laboratory.

### Cell Culture

Mouse embryonic fibroblast (MEFs) derived from *Notch1^fl/fl^; CreESR1* embryos (E13.5) which were dissected in sterile PBS, non-mesenchymal tissues were removed, and remaining tissue was digested with 0.25% trypsin (Wisent) at 37 °C for 10 min. Both MEFs and bovine aortic endothelial cells (BAECs; VEC Technologies, Rensselaer, NY, USA) were cultured in DMEM (Wisent) supplemented with 10% FBS (Gibco), 100 U/ml penicillin, 100 μg/ml streptomycin, and 50 μg/ml gentamicin. Cells were maintained at 37 °C in a humidified atmosphere of 5% CO₂ and 20% O₂, with medium replaced every 48 h, and passaged at ∼90% confluence using 0.25% trypsin.

### Tamoxifen injection of mice

To activate Cre recombinase, tamoxifen (4-hydroxytamoxifen; Sigma-Aldrich, T5648) was dissolved in corn oil (Sigma, C8267) at 1 mg/ml. *Notch1^fl/fl^; Cdh5-CreERT2* mice received 50 μl tamoxifen by intragastric injection at the milk spot from postnatal day 1 (P1) to postnatal day 3 (P3). Endothelial-specific Notch1-deficient mice are referred to as *Notch1^cKO^*, and tamoxifen-treated Cre-negative littermates were used as controls (*Notch1^cWT^*).

### Cell treatments

To induce Notch1 deletion *in vitro*, *Notch1^fl/fl^; CreESR1* MEFs were treated with 2 and 4 µM tamoxifen for four consecutive days, with medium refreshed every 48 h. Cre-positive MEFs without tamoxifen treatment served as controls. For pharmacological Notch1 inhibition, DAPT (Sigma-Aldrich) was dissolved in DMSO and applied at 20, 40, and 60 µM to wild-type MEFs and BAECs for 48 h.

### Proliferation assays

MEFs were seeded in 12-well plates and cultured for 72 h. Four fields per well (three wells per condition) were imaged with a 20× objective in three independent experiments. Cell growth rates were normalized to cell numbers at day 1 (0 h after adherence). Cell counting was performed using ImageJ software. Data represent the mean cell number ± standard error of the mean (SEM) from three wells at each time point.

### Migration assays

Cell migration was assessed using a scratch wound-healing assay. Confluent monolayers of MEFs and BAECs were cultured in 24-well tissue culture plates in complete growth medium. A linear wound was generated by scraping the monolayer with a sterile 200-µl pipette tip. Detached cells were removed by gently rinsing twice with PBS (1×), and cultures were replenished with fresh complete medium.

Time-lapse bright-field imaging was performed using a Nikon transmitted-light microscope (DS-Fi1 color camera). Images were acquired immediately after wounding (0 h) and after 8 h of incubation. Wound areas were quantified using ImageJ software by manually delineating the cell-free regions. Cell migration was calculated as the reduction in wound area between 0 h and 8 h. For each experimental condition, the mean wound closure from three replicate wells was determined and expressed relative to the control to obtain fold-change values.

### Gene set enrichment analyses (GSEA)

GSEA was conducted using GSEA v4.0.1 software provided by Broad Institute of Massachusetts Institute of Technology and Harvard University. We used GSEA to validate correlation between molecular signatures in phenotypes of interest. Data analysis was performed on the single-cell RNA-seq dataset published by Chen et al., IOVS 2023 [56]. Following the same pipeline described in the Methods, we reanalyzed the scRNA-seq data and used the FindMarkers function in Seurat to identify differentially expressed genes in P3 *Fz5^-/-^* mutant versus P3 *Fz5^+/+^* wild-type (WT) mice. The resulting ranked gene list was used as input for subsequent Gene set enrichment analyses (GSEA). Notch related genesets were obtained manually from the literature and from the Gene Expression Omnibus (GEO). Pathway-based gene sets were obtained from Reactome for enrichment analysis.

### Flow cytometry

Hyaloid vessels were isolated from postnatal mice at P0, P4, and P8, dissociated using 1× trypsin (MULTICELL, Cat. 325-542-CL) and washed with 1× PBS. For surface marker detection, cells were resuspended in PBS and incubated with PECAM-1 (CD31)-PE (Santa Cruz Biotechnology; sc-376764 PE) for 30 min on ice, followed by washing with PBS. Apoptotic cells were labeled with Annexin V-FITC (BioLegend; Cat. 640906) by incubating samples for 15 min at room temperature. All staining steps were performed protected from light using aluminum foil. Flow cytometric acquisition was performed on a BD Accuri C6 Plus (2-laser, 4-color), and data were analyzed using BD Accuri C6 Plus Analysis software.

### Collection of pre and postnatal eyes for immunofluorescence

Pregnant females were euthanized at embryonic day 18.5 (E18.5). Following a midline abdominal incision, uterine horns were exposed, and fetuses were removed from the amniotic sacs. Embryos at E18.5 and pups at postnatal day 0 (P0), P4, and P8 were sacrificed by decapitation. Eyes were enucleated, and a small puncture was made through the cornea using a 26G needle. Samples were fixed in 4% paraformaldehyde (PFA) for 1 h at room temperature, followed by three washes in PBS. Eyes were either processed immediately at room temperature or stored in PBS at 4 °C until further use.

### Immunofluorescence

For visualization of hyaloid and retinal vasculature in embryonic and postnatal mice, eyes were enucleated and fixed in 4% PFA at room temperature, followed by three washes in cold PBS. Hyaloid tissues were permeabilized and blocked in PBS containing 10% FBS, 1% Triton X-100, and 0.01 M glycine. Retinas were treated with a permeabilization-blocking solution composed of 3% BSA, 3% FBS, 0.01% sodium deoxycholate, and 0.5% Triton X-100 in PBS.

For blood vessels staining, hyaloids and retinas were incubated with isolectin B4 (IB4, 1:200; DL-1208-.5, CA, Newark) in the corresponding blocking solution. For protein localization, flat-mounted hyaloids and retinas were incubated with primary antibodies (see Table S1), followed by secondary antibodies: donkey anti-rabbit Alexa Fluor 488 IgG (1:500, A21206; Invitrogen) or donkey anti-rabbit Alexa Fluor 594 IgG (1:500, A21207; Invitrogen). After staining, tissues were washed in PBS and counterstained with DAPI (1 µg/ml, D9542; Sigma).

Retinas were flat-mounted directly, whereas hyaloids were embedded in 2% ultra-low-melting-point agarose (800-016-TC) and incubated at 37 °C in a humidified chamber to allow gradual gel diffusion. Hyaloids were then transferred to 4 °C and subsequently flat-mounted. All samples were mounted in glycerol (Sigma-Aldrich) and imaged using a confocal microscope (Olympus FV1000) with 10× and 60× oil-immersion objectives. Images were processed using Photoshop CS4 (Adobe Systems). Immunofluorescence quantification was performed on 3-5 Z-Stack images acquired at 60× of hyaloid vessels using ImageJ software.

### EdU labeling and Imaging

To label proliferating cells *in vivo*, the thymidine analog EdU (5-Ethynyl-2′-deoxyuridine, 50 µg/g body weight, in PBS) was injected into postnatal 8 (P8) mice. After 6 hours, the eyes were enucleated and fixed with 4% PFA for 1 hour at room temperature. EdU labeling was detected using the Click-iT^TM^ EdU Alexa Fluor 647 Imaging kit (Invitrogen, Cat: C10086).

### Quantification of hyaloid vascular parameters

Hyaloid vascular density, vessels diameter and vascular junctions in flatmounted hyaloids vessels was analyzed using ImageJ software as described before [70].

### RNA extraction and real-time quantitative PCR

Total RNA was extracted and suspended in commercial RNase-free water with a RNeasy Minikit (Qiagen, Germany). After DNase I treatment, cDNA was synthesized from 1 µg total RNA using the SuperScript II Reverse Transcriptase kit for RT-PCR (Thermo Fisher Scientific) according to the manufacturer’s instructions. cDNA (5 ng) was amplified in triplicate using SYBR Select Master Mix or Power Up SYBR Master Mix (Thermo Fisher Scientific).

Gene expression analysis was performed using the comparative cycle threshold (ΔCT) method, normalized with the expression of reference genes β-actin and presented as the mean fold change (± SEM) compared with control samples. The specific mouse primers pairs (Forward/Reverse) from the Invitrogen company are listed in Table S2.

### Western blot

Hyaloid vessels or cell samples were lysed in NP40 and protein content extracted using RIPA lysis buffer with mechanical help. After quantification, protein probes were loaded in polyacrylamide gels and separated using an SDS-page electrophoresis. Proteins were then transferred to a Nitrocellulose membrane and blocked in 3% skim milk in TBS-T for 1 h at room temperature. Membranes were incubated with diverse antibodies (Table S3) overnight at 4 C, followed by a 1 h incubation with species-appropriate HRP-conjugated secondary antibodies. Proteins were detected with ECL clarity method (Biorad) in a chemiluminescence imaging system.

### Statistical analysis

Data were presented as mean ± standard error of the mean (SEM) or as values. Statistical significance was assessed using grouped two-tailed Student’s t-tests. Images were quantified with ImageJ software, and statistical analyses were performed in GraphPad Prism 9.0 (GraphPad Software, San Diego, CA). It was considered statistically significant if the *p* value was less than 0.05 (*), 0.01 (**), 0.001 (***) or 0.0001 (****). N is indicated pertinently for each experiment.

## Supplementary Materials

**Table S1:** Primary antibodies used for Immunofluorescence.

| Antibody | Species | Clone | Dilution | Company | Catalog # |
| --- | --- | --- | --- | --- | --- |
| Ki-67 | Rabbit | monoclonal | 1:300 | ABclonal | A11390 |
| C-CASP3 | Rabbit | monoclonal | 1:300 | ABclonal | A19654 |
| IBA1 | Rabbit | monoclonal | 1:300 | ABclonal | A19776 |
| ERG | Rabbit | monoclonal | 1:500 | CellSignaling | A7L1G |
| NG2 | Rabbit | monoclonal | 1:300 | CellSignaling | 43916 |
| α-SMA | Rabbit | monoclonal | 1:300 | CellSignaling | 19245 |

**Table S2:**
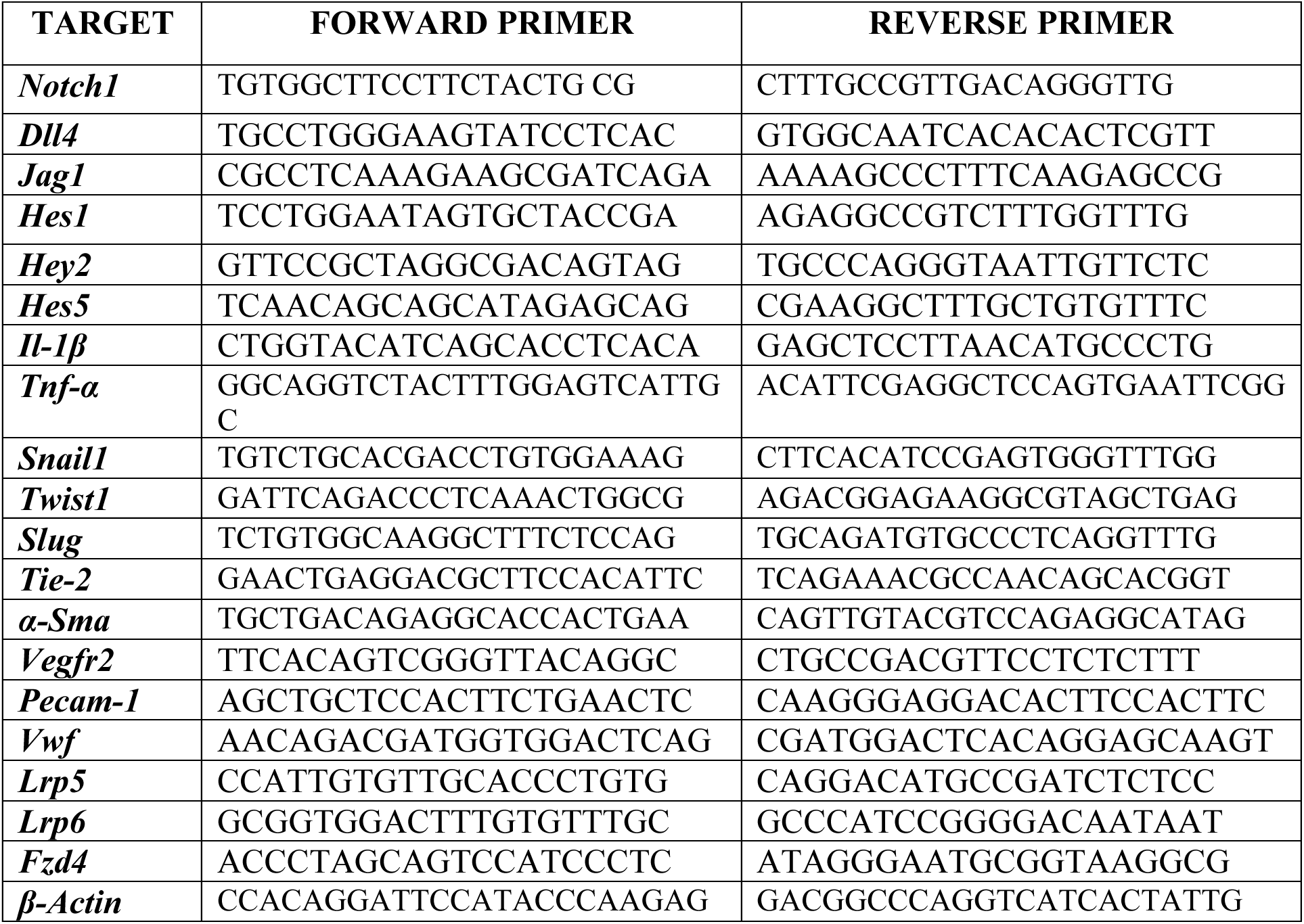
Oligonucleotides used for qRT-PCR reagents of murine origin.

**Table S3.** Primary antibodies used for Western blot analysis.

| Antibody | Species | Clone | Dilution | Company | Catalog # |
| --- | --- | --- | --- | --- | --- |
| β-actin | Mouse | monoclonal | 1:10000 | ABclonal | AC004 |
| Notch1 | Rabbit | polyclonal | 1:1000 | ABclonal | A7636 |
| NICD | Rabbit | polyclonal | 1:500 | ABclonal | A22674 |
| DLL4 | Rabbit | polyclonal | 1:1000 | ABclonal | A12943 |
| JAG1 | Rabbit | polyclonal | 1:1000 | ABclonal | A12733 |
| HES1 | Rabbit | monoclonal | 1:500 | ABclonal | A0925 |
| HEY1 | Rabbit | polyclonal | 1:500 | ABclonal | A16110 |
| CASP3 | Rabbit | monoclonal | 1:1000 | ABclonal | A19654 |

## Supplementary figures

**Fig. S1.**
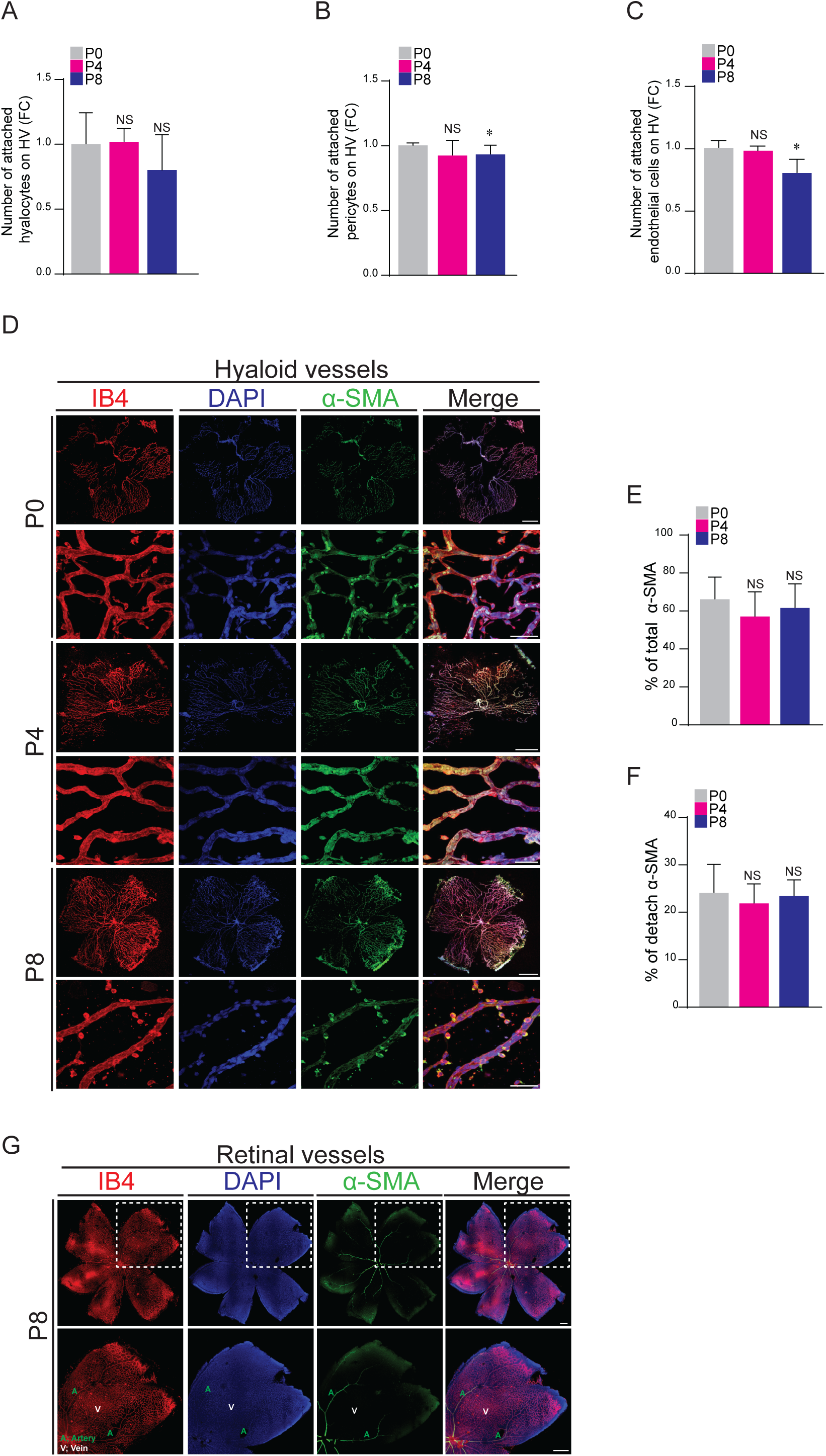
Hyaloid vessel-associated immune and mural cell dynamics postnatally. (A) Quantification of attached IBA1^positive^ hyalocytes, (B) NG2^positive^ pericytes and (C) ERG1^positive^ endothelial cells on P0, P4, and P8 hyaloid vessels (HV) (n = 3). Results are expressed as fold change (FC) normalized to P0 stage. (D) Representative confocal micrographs of P0, P4, and P8 hyaloid vessels flatmounts labeled with vascular smooth muscle cells marker (α-SMA), IB4 and DAPI. (E, F) Quantification of the percentage of total and detached α-SMA show constant expression during hyaloid vascular regression (*n* = 3). (G) Representative confocal immunofluorescence staining of α-SMA, IB4 and DAPI of flatmounted P8 retinal vasculature reveals well-functioning immunostaining of α-SMA in arteries but not in veins (n = 3). Data are means ± SEM. Represented p-values are *≤0.05 from unpaired T-test. Non-significant (NS). Scale bars, 100 µm and 200 µm [for higher-magnification images in (D, G)].

**Fig. S2.**
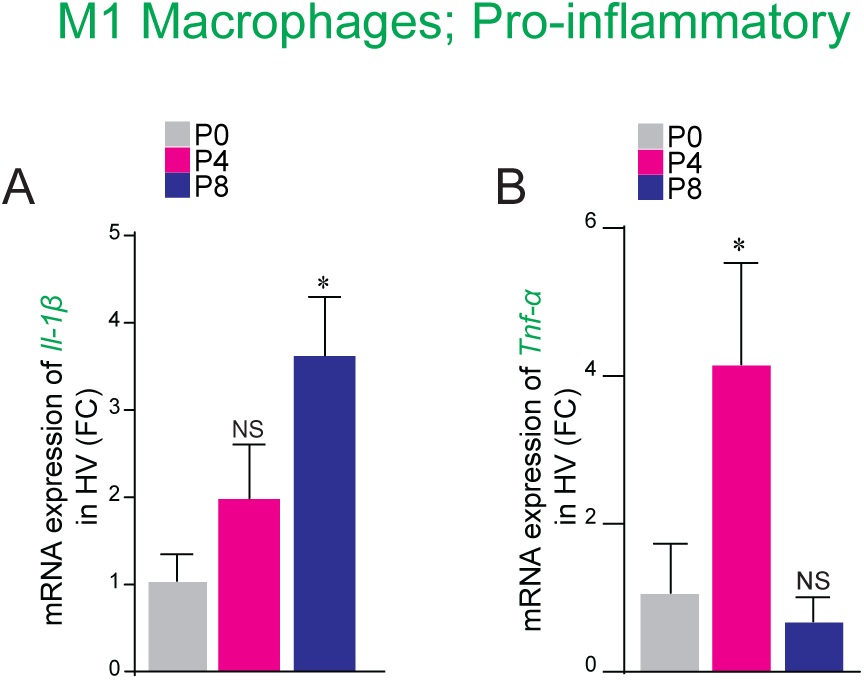
Endothelial detachment during hyaloid regression is associated with pro-inflammatory signaling. (A) qRT-PCR shows induction of expression of pro-inflammatory transcripts *I1-1β* and (B) *Tnf-α* at P8 and P4 hyaloid vessels (HV) stages respectively versus P0 stage. β-actin was used as a reference gene. Results are expressed in fold change (FC) compared to P0. Data are means ± SEM. Represented p-values are *≤0.05 from unpaired T-test. n = 3 for *Il-1β* and *Tnf-α* (A-B). Non-significant (NS).

**Fig. S3.**
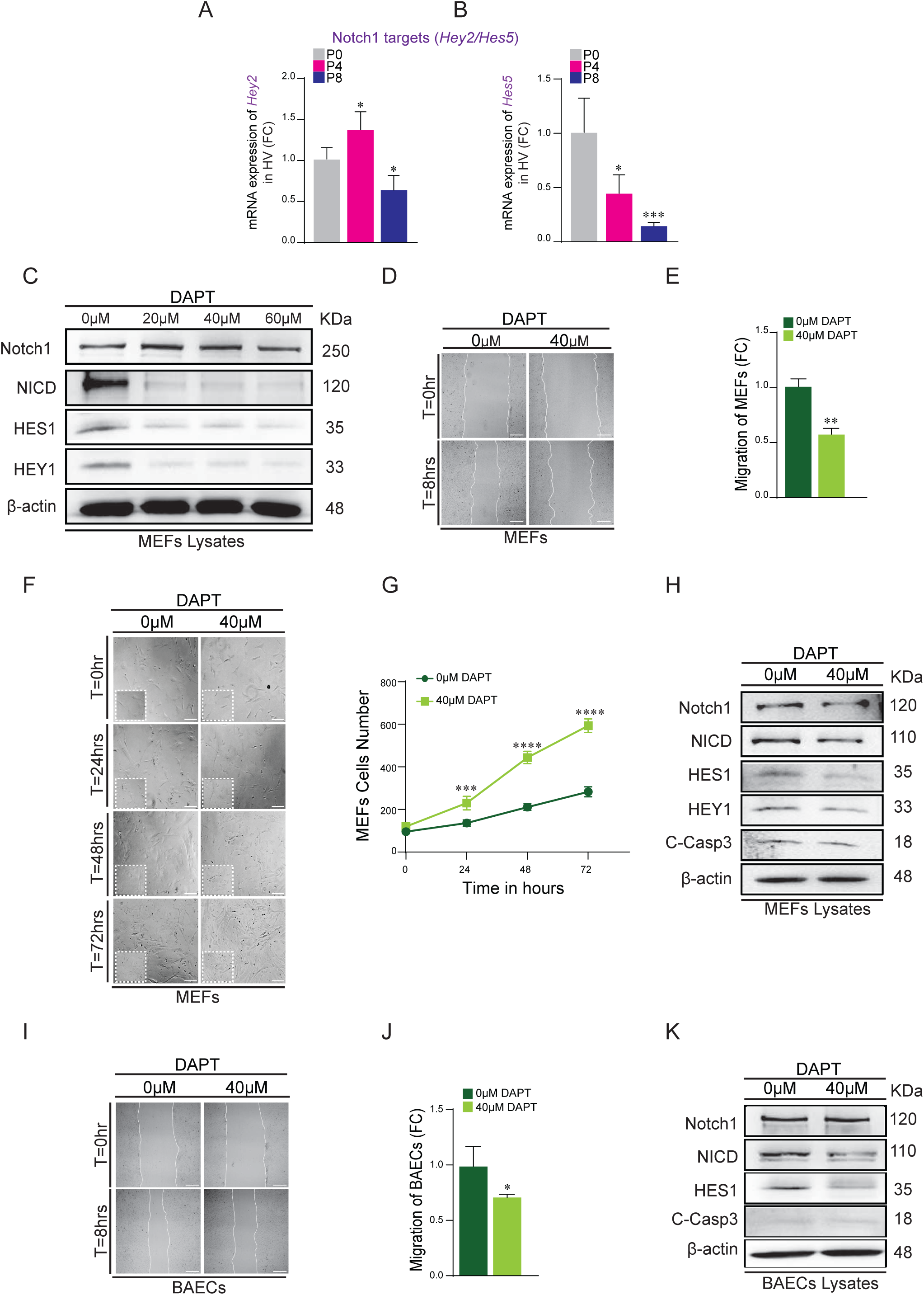
Notch1 inhibition decreases cell proliferation and migration *in vitro*. (A) qRT-PCR for levels of Notch1 target genes *Hey2* and (B) *Hes5* measured in hyaloid vessels (HV) at P4 and P8 versus P0. β-actin was used as a reference gene. (C) Western blot analysis of Notch1, its active form (Notch1 intracellular domain, N1ICD), HES1 and HEY1 protein expression levels in mouse embryonic fibroblasts (MEFs). Cell lysates from MEFs treated with DAPT in a dose-dependent manner of 20, 40, and 60µM for 2 days. β-actin is used as a loading control. (D) Representative images from the wound edge at the initiation of imaging (0 hours) and 8 hours after wounding of MEFs monolayers, treated with 40 µM DAPT show a decrease in migration compared to non-DAPT treated control, and quantified in (E). (F) Representative images of MEFs from the time-course proliferation assay at the initial time-point (0 hours) and after 24, 48 and 72 hours, treated and non-treated with 40µM DAPT, show increased growth with DAPT treatment, quantified in (G). (H) Immunoblot analysis of Notch1, N1ICD, HES1, HEY1, and C-Casp3 protein expression levels in MEFs lysates treated with 40µM DAPT compared to non-treated MEFs control. β-actin is used as a loading control. *n =* 3 experiments. (I) Representative images from the wound edge at the initiation of imaging (0 hours) and 8 hours after wounding of BAECs monolayers, treated with 40 µM DAPT show a decrease in migration compared to non-DAPT treated BAECs, and quantified in (J). (K) Immunoblot analysis for Notch1, N1ICD, HES1, and C-Casp3 of BAECs lysates treated with DAPT versus non-treated control. β-actin was used as a loading control. *n* = 3 experiments. Data are means ± SEM. Represented p-values are *≤0.05, **≤0.01, ***≤0.001, ****≤0.0001 from one-way ANOVA with unpaired T-test. n = 4-5 biological samples for *Hey2* (A); n = 3-5 biological samples for *Hes5* (B). *n* = 3 biologically independent experiments of proliferation (F, G) and migration (I, J). Non-significant (NS); Results are expressed in fold change (FC) compared to non-treated MEFs and BAECs. Scale bars, 100 µm (D, E); 200 µm (F).

**Fig. S4.**
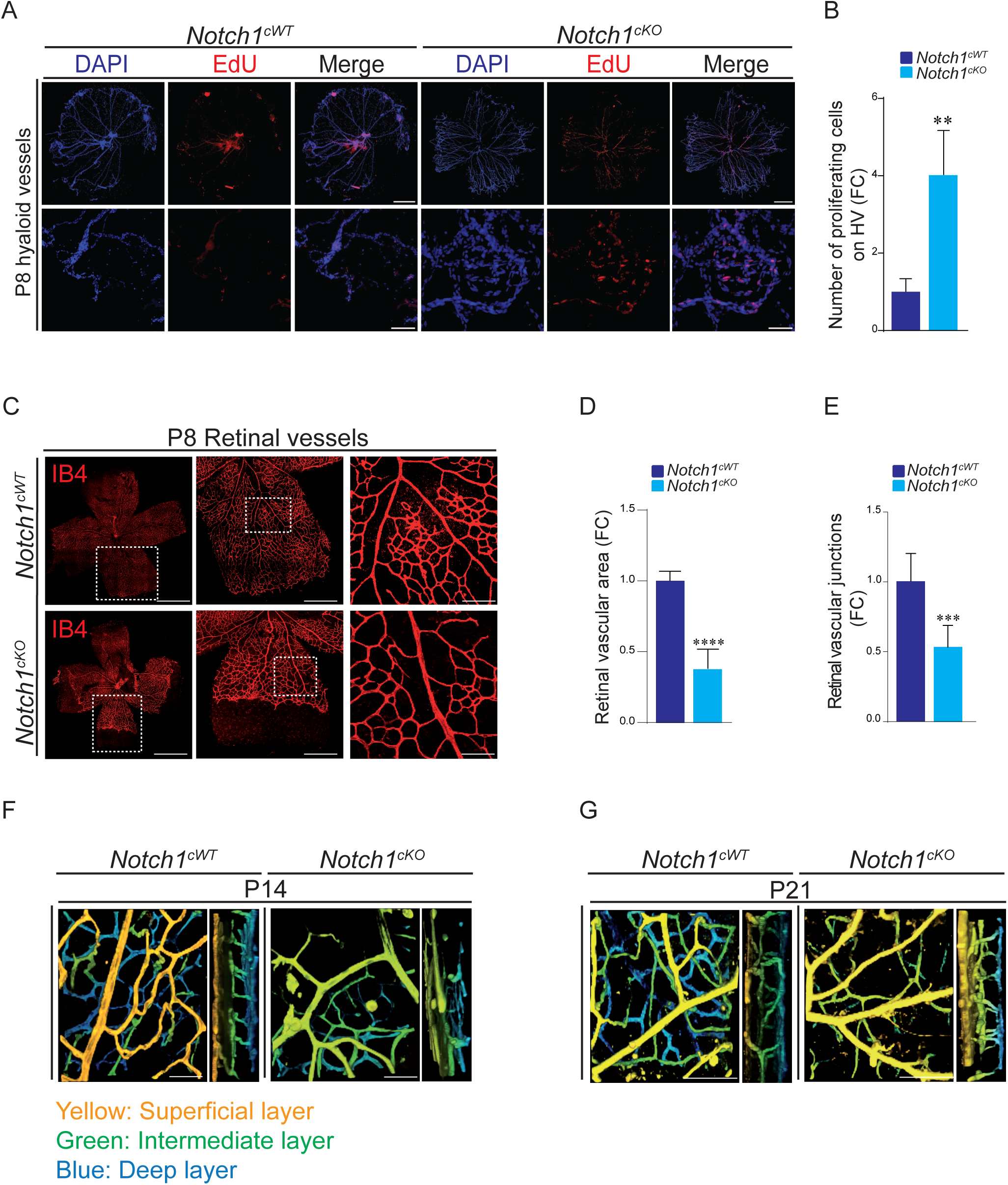
Endothelial-specific Notch1 deletion leads to retardation in retinal vascular superficial layer formation. (A) Representative confocal micrographs of P8 hyaloid vessels (HV) flatmounts immunostained with EdU and DAPI of *Notch1^cKO^* versus *Notch1^cWT^* and the number of proliferative (EdU positive) cells quantified in (B) (*n* = 3). Results are expressed as fold change (FC) compared to *Notch1^cWT^*. (C) Representative IB4 staining and (D) quantification of retinal vascular area, and (E) vascular junctions in P8 flatmounted retinal vasculature in *Notch1^cKO^*versus *Notch1^cWT^* (*n* = 7). (F, G) Three-dimensional (3D) P14 and P21 retinal vasculature shows 3 vascular layers [first superficial layer (yellow), second deep layer (blue) and second intermediate layer (green)] in *Notch1^cKO^*versus *Notch1^cWT^* (*n* = 3). Data are means ± SEM. Represented p-values are **≤ 0.01, ***≤ 0.001, ****≤0.0001. Scale bar, 100 µm (A), 200 µm [for higher magnification images in (A, C, F, G)].

**Fig. S5.**
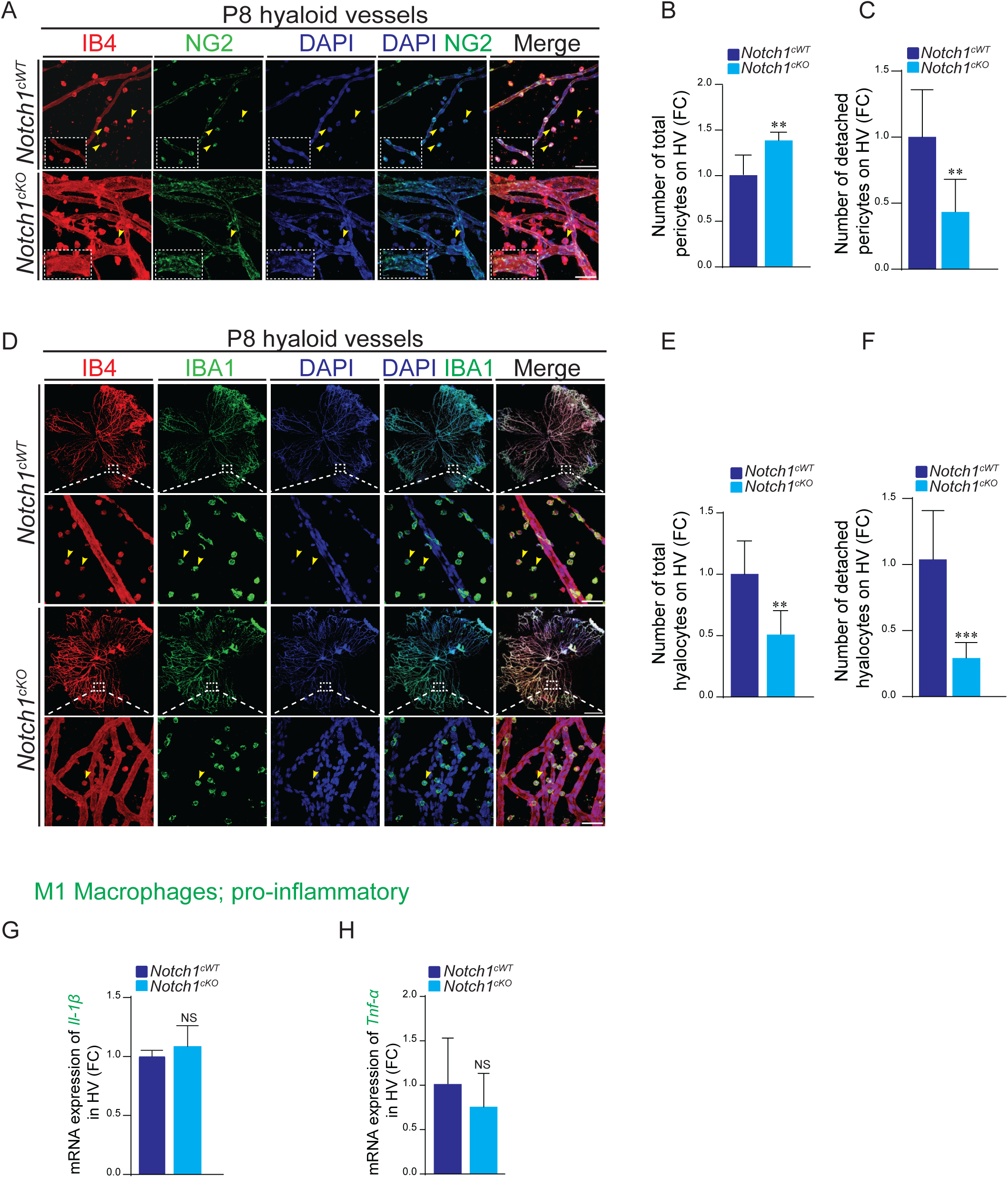
Loss of endothelial Notch1 alters hyaloid pericyte positioning and macrophage activation. (A) Representative confocal micrographs of P8 hyaloid vessels (HV) flatmounts immunostained with NG2, IB4 and DAPI in *Notch1^cKO^* versus *Notch1^cWT^*. (B) Quantification of total and (C) detached (yellow arrowheads) NG2^positive^ pericytes in hyaloid vessels of *Notch1^cKO^* compared to *Notch1^cWT^* mice (*n* = 3). (D) Representative IBA1, IB4 and DAPI staining of P8 *Notch1^cKO^* versus *Notch1^cWT^* hyaloid vessels. (E) Quantification of total and (F) detached (yellow arrowheads) IBA1^positive^ hyalocytes on hyaloid vessels of *Notch1^cKO^* compared to *Notch1^cWT^* mice (*n* = 3). (G) qRT-PCR shows constant transcript levels of *Il-1β* and (H) *Tnf-α* in hyaloid vasculature of *Notch1^cKO^* versus *Notch1^cWT^*. Data are means ± SEM. Represented p-values are **≤0.01, ***≤0.001 from unpaired T-test. *n* = 4 for *Il-1β* (G) and *n* = 3 for *Tnf-α* (H). Non-significant (NS). Results are expressed as fold change (FC) normalized to control *Notch1^cWT^*.

